# FORUM: Building a Knowledge Graph from public databases and scientific literature to extract associations between chemicals and diseases

**DOI:** 10.1101/2021.02.12.430944

**Authors:** M. Delmas, O. Filangi, N. Paulhe, F. Vinson, C. Duperier, W. Garrier, P.-E. Saunier, Y. Pitarch, F. Jourdan, F. Giacomoni, C. Frainay

## Abstract

Metabolomics studies aim at reporting a metabolic signature (list of metabolites) related to a particular experimental condition. These signatures are instrumental in the identification of biomarkers or classification of individuals, however their biological and physiological interpretation remains a challenge. Overcoming this challenge is critical when aiming to associate metabolic signatures with potential pathological outcomes. To support this task, we introduce FORUM: a Knowledge Graph (KG) providing a semantic representation of relations between chemicals and biomedical concepts, built from a federation of life science databases and scientific literature repositories. An important number of scientific articles discuss relations between chemical compounds and biomedical concepts in various contexts, from biomarkers to therapeutic uses. The extraction of these statements and their interconnection in a graph structure can thus allow us to identify and explore relations strongly supported in the scientific literature.

The use of a Semantic Web framework on biological data allows us to apply ontological based reasoning to infer new relations between entities. We show that these new relations provide different levels of abstraction and could open the path to new hypotheses. We estimate the statistical relevance of each extracted relation, explicit or inferred, using an enrichment analysis, and instantiate them as new knowledge in the KG to support results interpretation/further inquiries. Beyond this result, FORUM can also provide insights into complex biological questions and the extracted information could then be used for further developments.

Containing more than 8 billion triples and providing more than 8 million relations, FORUM leverages the increasing availability of linked datasets in life science and is built in agreement with FAIR principles. A web interface to browse and download the extracted relations, as well as a SPARQL endpoint to directly probe the whole FORUM knowledge graph, are available at https://forum-webapp.semantic-metabolomics.fr. The code needed to reproduce the triplestore is available at https://github.com/eMetaboHUB/Forum-DiseasesChem.

## 1 Introduction

Among all omics sciences, metabolomics stands the closest to phenotype as it captures at the highest level the response of the system to environmental changes, physiological disorders or stimulus [1]. The profiling of small molecules in a tissue, a fluid or an organism using high throughput untargeted analysis is a powerful tool to identify biomarkers [2]. It offers valuable insights for diagnosis and patient prognosis, and opens promising opportunities for drug design and physiopathology understanding. However, the current challenge of metabolomics is to exploit these metabolic signatures to their full extent by connecting the metabolic disruptions to the observed phenotypes, giving insights into the underlying mechanisms [3]. While models of metabolism become more accurate, yielding valuable insight from metabolic network analysis, this challenging issue requires knowledge beyond biochemical transformations alone, connecting metabolites to higher-order phenotypical information.

Scientific literature, as a primary way of reporting and sharing scientific progress, is an obvious valuable resource for interpretation of metabolic profiles. However, the amount of information that needs to be processed in order to meet interpretation requirements is at the core of the current limitations. This calls for computational support to deal with the information overload, which requires modelling knowledge in an appropriate form. With digitalization, annotation and indexing of scientific literature, information can be automatically extracted, paving the way for more machine-readable knowledge representation. In addition, federation of structured data from various sources to create an exploitable centralized knowledge base could empower large-scale assisted reasoning, and thus constitute an interesting step toward closing the gap in metabolomics.

Storing information and modeling knowledge as networks have been proven useful in various domains[4]. The resulting Knowledge Graphs (KG) are especially suited for information retrieval tasks that require bridging information through indirect relationships in complex networks. A KG is a formal and structured representation of knowledge, as a graph of entities and relations, commonly described in a triple formalism (Subject-Predicate-Object). In a KG, each entity, property or relation, could also be described by a semantic layer, precisely defining the related concepts using structured and standardized vocabularies, usually from the appropriate ontologies [5]. The last decade has seen the rise of available representation of knowledge in such formalism, highlighting the potential of Semantic Web technology to aggregate and exploit these datasets [6, 7]. The most important data providers in life sciences propose the content of their databases in RDF format [8, 9], a standard of the Semantic Web, particularly suited for building KG.

One of the most used repositories of knowledge is the PubMed database that hosts more than 30 million scientific articles from life science. While articles hold relevant information to make sense of metabolomic data, extracting it from unstructured text is a complex issue[10]. Fortunately, these articles are manually annotated by *National Library of Medecine* (NLM) trained experts, using MeSH descriptors (Medical Subject Headings), to describe their main topics. The MeSH Thesaurus^1^ is a controlled vocabulary containing a broad range of biomedical concepts, hierarchically organised with parent-child relationships, from generic descriptors (eg. *Cardiovascular Diseases*) up to representatives of specific concepts (eg. *Coronary Occlusion*). This metadata is primarily used for information retrieval purpose, fueling the PubMed search engine. Being computationally readable, unambiguous and less noisy than text, as well as also being accessible as RDF, several resources exploit them for supporting data interpretation by performing statistical analysis of corpora’s MeSH distribution [12, 13]

Beyond the recent surge in data availability, a tremendous effort has also been put into the interoperability of life-science resources, by extensive cross-references between databases and matching tools (eg. UniChem [14], MetaNetX [15] and *E-utilities*^2^ *and BridgeDb[16])*. Among these databases lies PubChem [17], the world’s largest free chemistry database, providing datasets that gather data and properties associated with chemical compounds [8], including mentioning articles in PubMed[18]. By navigating the cross references between resources like Pub-Chem and PubMed, it is thus possible to link compounds to biomedical concepts. Moreover, these two entities are semantically described by ontologies or thesauri, with definitions, synonyms and other metadata, increasing inter-operability and data integration [5]. Leveraging the Semantic Web framework, relationships between concepts in the MeSH hierarchy can also be taken into account. The same can be achieved by integrating semantic annotation of compounds, using chemical classification ontologies such as ChEBI[19] and ChemOnt[20]. Those relationships enable some degrees of abstraction allowing, for instance, reasoning over chemical families rather than individual compounds, raising new hypotheses through in-class generalization. Moreover, this abstraction can be implicitly performed by the KG’s reasoner using the *true-path rule*. Basically, the *true-path rule* states that if an entity is annotated to a class, it implicitly also belongs to its parent classes. This rule is also used in the context of information retrieval: the Pubmed search engine uses this property to retrieve, for example, articles indexed with the MeSH descriptor “Parkinson’s disease” from a query targeting articles about the broader concept of “Neurodegenerative Diseases” [21].

Finally, all the content of these resources and their semantic descriptions can be combined to build a Knowledge Graph, a network of semantically-described entities connected by factual relationships, that hosts the information required to support knowledge discovery for metabolomics.

Some tools have already been developed to extract and propose gene or drug candidates related to pathological phenotypes based on literature evidence and databases [22, 23, 24, 7]. In a context closer to metabolomics, some approaches have also been developed to explore relations between chemicals and biomedical concepts based on the scientific literature, and most of them use text-mining approaches such as PolySearch [25], Alkemio [26], LimTox [27] or RELigator [28]. These approaches have a fair sensibility and can identify close relations between terms from co-mentions in single sentences. However, they usually offer a low precision, requiring each association be checked manually, through analysis of the sentences that support it and usually the whole articles that contain them. The KG being a resource built to infer new hypotheses from large-scale existing associations, those constituting it must be integrated to the lowest false-positive rate, beyond current text-mining tools performances. Indeed, while a missing link would not necessarily obfuscate a latent relationship, thanks to path redundancy in the KG, a spurious link between unrelated concepts could substantially alter the KG structure. The use of the NLM indexation system with the MeSH vocabulary can help to avoid these issues by leveraging manual labelling performed by trained experts. The association at the article topic level also grasps the full context and helps to avoid the ‘noise’ and ambiguities found at the sentence level.

Another tool has taken advantage of the linking of MeSH terms to PubChem compounds: Metab2MeSh [29] was a web server dedicated to the annotation of individual compounds with MeSH descriptors. It was a valuable resource that is unfortunately no longer available at the date of writing. Metab2MeSH aimed at testing the statistical significance of co-occurring concepts and compounds in literature, based on Over-Representation Analysis (ORA). ORA is a widely used approach to bring biological meaning to results, like Pathway enrichment [30] or Gene Set Enrichment Analysis (GSEA) [31], and is commonly performed on sets tested independently, setting aside relationships among them. Metab2MeSH is based on a similar methodology, effectively ignoring the MeSH thesaurus structure. Consequently, it considers “thematic sets” of articles distinct from what a user would obtain by querying such a topic in PubMed, as well as missing higher-level associations that could be inferred. Nevertheless, some ORA methods account for relations between entities. For instance, GO Enrichment Analysis[32] is a well used resource to compute functional enrichments on gene sets, where those related to GO-terms are determined by considering their hierarchical relations in the Gene Ontology [33] using the *true-path rule* [34]. Applying this approach on the MeSH thesaurus or chemical ontologies can thus have a significant impact on predicted relations by the ORA analysis.

Despite related end goals, none of the cited methods aims at providing a queryable Knowledge Graph to build upon, and they also lack open source implementation. The call for compliance to FAIR principle *(Findable, Accessible, Interoperable* and *Reusable*) has recently grown strong, as it improves the sharing and exploitation of an increasing amount of available data [35, 36]. Standing at the crossroad of multiple disciplines, current challenges in metabolomics need to be answered with integrative infrastructures, federerating knowledge, data and assertions, that should be managed respecting FAIR principles [37].

In this way, we have developed FORUM, an open and fully accessible Knowledge Graph, which allows the navigation of a network of relations between molecules and biomedical concepts by connecting PubChem, MeSH, ChEBI, ChemOnt and MetaNetX. We provide some examples to illustrate what kind of relations can be extracted from FORUM and see how their connections inside the KG can help to explore new hypotheses. Beyond its support for results interpretation, this is a proof of concept meant to illustrate the potential of open linked data in metabolomics, as FORUM could be easily extended for various purposes and grow with the endorsement of FAIR principles by the metabolomics community.

FORUM currently holds associations between 344.769 PubChem compounds, 6.740 ChEBI classes and 2.783 ChemOnt classes to 24.382 MeSH descriptors, implying more than 8.7 million of significant associations, with 1.096.106 related to diseases.

All associations are queryable through the web-portal at https://forum-webapp.semantic-metabolomics.fr. Data are accessible by connecting on the sftp server at ftp.semantic-metabolomics.org (see information on webportal) and ready-to-mine using the Enpoint at https://forum.semantic-metabolomics.fr, also allowing to exploit the connectivity between entities in custom requests to explore more complex questions.

## 2 Method

### 2.1 Integrated data resources

The main goal of FORUM, our developed KG, is to provide links between chemical compounds and biomedical concepts supported by literature. It is composed of several inter-connected sub-graphs, each describing particular entities and relations (Figure 1A). Each entity is identified using a specific and persistent *URI* (Uniform Resource Identifier) and relations between them are described by predicates in RDF triples format. An example of this formalism is presented in Figure 1B, showing some relations between compounds, literature and chemical classes that can be extended from the Glucose record (compound:CID5793) in the KG.

**Figure 1:**
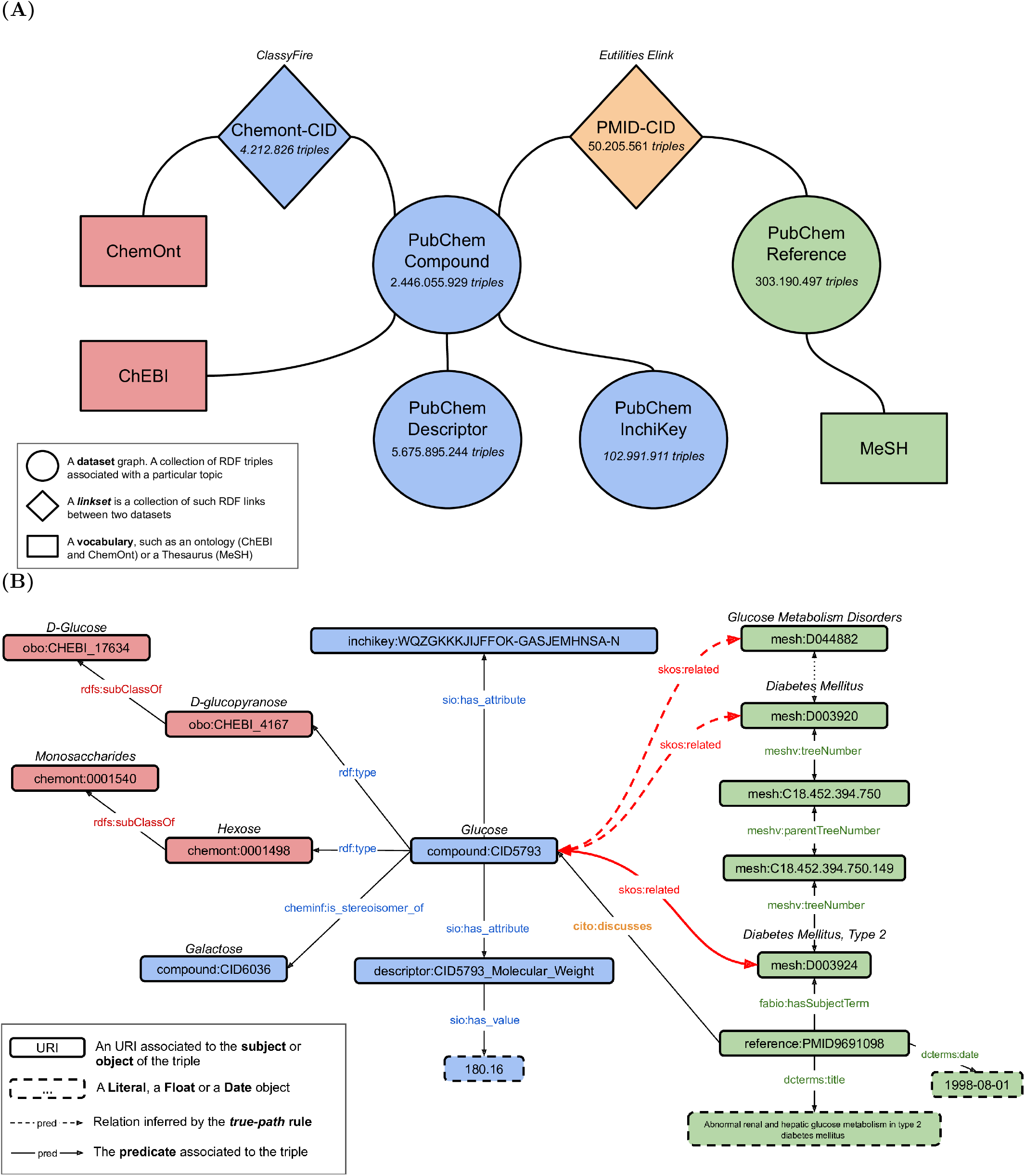
A general view of the Knowledge Graph architecture (**A**) and an example of interrelations between compounds, literature and chemical classes, that can be extracted from the knowledge graph, centered around Glucose (**B**). In (**A**), only graphs containing the basic relations between entities are indicated while graphs containing inferred relations from statistical analysis (*skos:related* relations) are hidden. Blue boxes represent triples exposing properties associated with PubChem compounds, green triples described publications, orange triples established links between molecules and publications, and red are related to chemical classification of molecules. In (**B**), grey arrows represent explicit relations that are instantiated in the original dataset and bold red arrows represent added relations, inferred from the enrichment analysis.

Chemical compounds are described in the PubChem RDF compound graph. It contains more than 103 million unique chemical structures extracted from PubChem Substance. Their properties, such as mass, IUPAC name and InChiKey identifiers are exposed in the PubChem Descriptor and PubChem InchiKey graphs (triples in blue on Figure 1B). In order to provide a semantic representation of chemicals, which allows to infer new relationships [5], we use the ChEBI^3^ ontology classification available for 116.068 pubchem compounds, provided in the PubChem store. We also integrated the ChemOnt^4^ ontology for 319.189 compounds using the web-services of ClassyFire[20] and the InChiKey annotation of compounds. The ChemOnt ontology file, originally formatted in OBO (Open Biomedical Ontologies), was converted to RDF syntax using Protege^5^ (v.5.50), to be imported in the KG. In ontologies, hierarchical dependencies between classes are defined using the predicate *rdfs:subClassOf*, as illustrated in the Figure 1B, where the Chemont chemical class of *Hexoses* is defined as a sub-class of *Monosaccharides*. PubChem also provides a representation of the Pubmed literature [8] associated with PubChem Compounds, Substances or Assays, in the PubChem Reference graph, including the MeSH indexation of each publication (triples in green on Figure 1B). All data extracted from PubChem RDF can be found on the NCBI ftp server^6^.

In order to build an explicit network of compound to article relations, we used the NCBI-Eutils Elink utility^7^, allowing the extraction of crosslinks between entities from different Entrez databases to define these links. Relations between PubChem compounds and Pubmed articles are provided from complementary sources: by external data contributors (eg. IBM Almaden Research Center^8^), by journal publishers or by matching PubChem chemical names on MeSH annotated to articles [18]. These links are then instantiated in a RDF graph, named *PMID-CID* (Figure 1A), with the *cito:discusses* predicate (in orange on Figure 1B), the CiTO ontology being particularly suited to describe bibliographic resources [38]. Contributors of each association are also referenced in an alternative graph, named *PMID-CID-endpoint*. The connections between compounds and articles then allow us to extract compounds to MeSH links from the chain of relationships embedded in our federation of resources. Those relations are symmetric, thus allowing to retrieve compounds from a MeSH descriptor as well.

The RDF graph of MeSH Thesaurus^9^ [39] provides a machine-readable representation of MeSH concepts, organized as a set of trees, where the *treeNumber* of a MeSH indicates its tree location. Then, the hierarchical relationships between descriptors in the MeSH tree can be accessed through the *meshv:parentTreeNumber* predicate (Figure 1B). Finally, additional ontologies were also integrated to define the nature of each entity or property used in the KG. For example, CiTO and FaBiO [38] describe bibliographic resources, Cheminf [40] defines chemical properties such as *is stereoisomer of*, dcterms^10^ references metadata, and SKOS^11^ is used to instanciate extracted relations.

The Knowledge Graph was built on a Virtuoso triplestore (version 7.20.3229) packaged in a Docker image^12^, providing access to data through the SPARQL endpoint. The SPARQL request language supports complex queries and allows to exploit the semantics defined in ontologies to extract stated relations and infer new ones from the KG. It also ensures the accessibility and the use of data and metadata available in the KG, following *FAIR* principles. The total number of articles associated with each compound, chemical class and MeSH descriptor, as well as the co-occurrences between them, were then determined using specific SPARQL requests against the endpoint. For associations with chemical classes, we only considered those with less than one thousand associated compounds, as classes like *Carboxylic acids and derivatives* may be too broad to bring meaningful results.

### 2.2 Computation of associations

By performing co-occurrence counts using the Virtuoso reasoner, associations between chemical entities or MeSH descriptors and PubMed articles are implicitly propagated to their ancestors in their respective ontologies/thesaurus, through the *subClassOf/parentTreeNumber* properties, according to the *true-path rule*. For example, this implies that, for a publication which is related to both *Glucose* (compound:CID5793) and *Diabetes Mellitus Type 2* (mesh:D003924), this article is also implicitly supporting the association between the chemical class of *Monosaccharides* (chemont:0001540) and the disease family of *Glucose Metabolism Disorders* (mesh:D044882), even if this relation was not explicit in the KG, Figure 1B. Associations were only computed on a selection of MeSH categories relevant in the studied context[29]: Diseases, Anatomy, Chemicals and Drugs, Phenomena and Processes, Organisms, Psychiatry and Psychology, Anthropology, Education, Sociology and Social Phenomena, Technology, Industry, and Agriculture.

The over-representation of each association in the scientific literature was then tested using a right-tailed fisher exact test, reporting *odds-ratio* and *p-value*, which was adjusted for multiple comparisons using the Benjamini-Hochberg procedure, proving *q-value*. Association can be ranked by effect-size using odds-ratios, and, to provide a ranking between associations using the q-value, even if it wasn’t precisely computable and was approximated to 0, the *χ*^2^ statistics is also indicated [29]. Associations with a *q-value* < 1*e* −6 are considered significant and are instantiated in specific assertion sub-graphs as follows: *chemical entity skos:related mesh:descriptor*, the *skos:related* predicate being a symmetric and non-transitive property used to assert associative links. An example is provided in Figure 1B, the instantiated relation between *Glucose* and *Diabetes Mellitus, Type 2* is assumed to be supported by other publications and is statistically significant. According to the *true-path* rule, the relation between Glucose and ancestors of *Diabetes Mellitus, Type 2* can also be inferred and instantiated. For clarity reasons, only these relations are indicated on the Figure, but in the same way, a *skos:related* relation could also exists between *Glucose* related chemical classes (eg. *Monosaccharides*) and *Diabetes Mellitus, Type 2*, as well as its parent descriptors.

The use of a threshold on the *q-value* is a convenient and widely used statistical approach to select relevant results from hypothesis tests, but is not sufficient for interpreting association strength [41]. Thus, we added the absolute number of supporting articles, as well as the creation of a complementary index named *fragility index*. We define the *fragility index* as the minimal number of articles, included in Jeffrey’s proportion confidence interval[42], that, when removed from the association corpus, increase the p-value over the significance threshold (1*e*− 6). Details of this approach are presented in supplementary materials (S3.2.1).

Code and SPARQL requests required to re-build all the Virtuoso triplestore and reproduce results are available at https://github.com/eMetaboHUB/Forum-DiseasesChem.

## 3 Results

### 3.1 Advantages of the semantic level on association extraction

The true-path rule affects corpora sizes of chemical classes and MeSH descriptors by propagating annotated literature to broader concepts and thus also influences counts used for statistical testing of independence. For instance, a broad descriptor like *Neoplasm* (mesh:D009369) goes from 96.373 explicitly annotated articles to 955.848 (*∼*10 fold) using literature related to sub-concepts. For more information, see Supplementary materials (S3.1). Beyond changes in counts, we analysed the consequence that the true-path rule induced on extracted relations. A Venn diagram of significant associations between MeSH descriptors and PubChem compounds detected with and without the use of the true-path rule is presented in Figure 2. The number of significant associations detected using propagation is more than twice of what is initially detected. More than half of these new associations link compounds to MeSH terms that are never directly co-annotated in the same article, but inferred from semantic relationships. Furthermore, 6% of initially detected associations were discarded by performing the analysis with the semantic elevation, as they no longer meet our inclusion criteria (*q-value* < 1*e* − 6).

**Figure 2:**
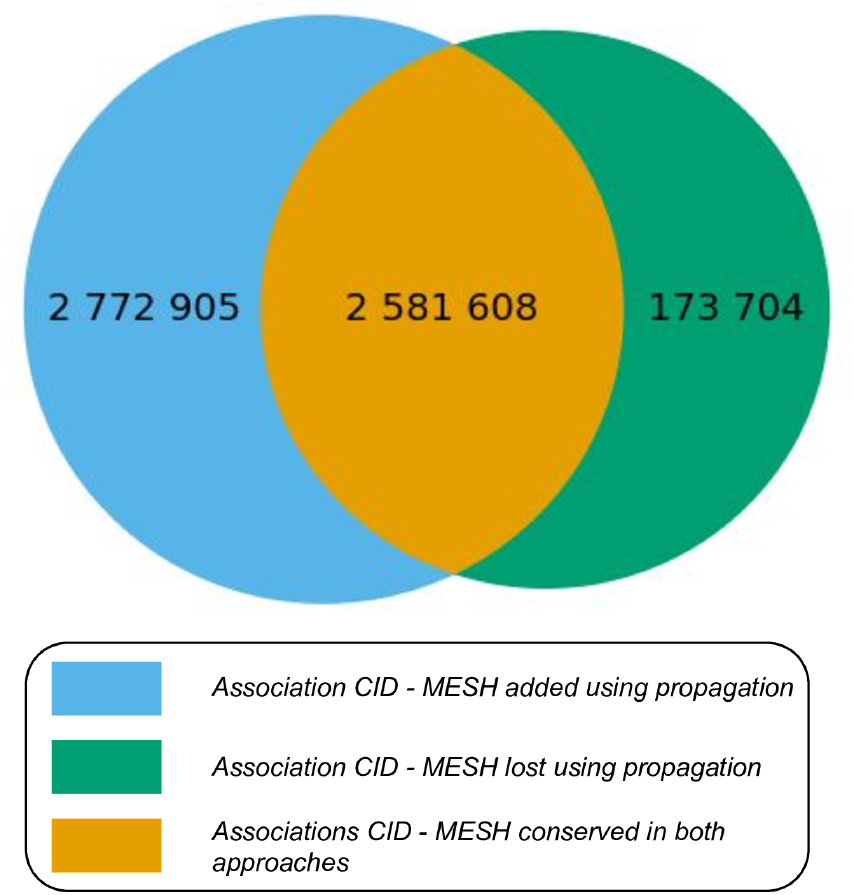
Venn diagram illustrating the overlap between sets of associations statistically significant (*q-value* < 1*e*−6) detected using or not using the literature propagation through the MeSH Thesaurus. Proportions are derived from log-transformation of sets’ size. 58% (1.614.120/2.772.905) of newly detected associations are completely novel as there are no direct links between these entities in the KG without using propagation.

To illustrate what kind of extracted associations can be affected by the method, we selected some typical examples and analyzed the MeSH annotation frequency of their supporting corpus, in order to illustrate the context surrounding the association. All MeSH descriptors explicitly annotated to publications supporting the co-occurrence between both entities were extracted and we selected the top 20 most important descriptors, using a score analogous to TF-IDF (See details in Supplementary materials). A low importance score indicates that the number of times a MeSH appears in the subset of articles describing the relation is close to what could be expected in a random subset of articles.

#### 3.1.1 Associations discarded by corpus size adjustment

The case of *Water* and *Eukaryote* describes the removal of associations due to the true-path rule. *Water* (CID 962) has a very large corpus of 156.845 annotated articles and an initial co-occurrence with *Eukaryote* of 330 publications. However, this appears highly significant (*p − value ≈* 2.9*e*− 49) as *Eukaryote* was explicitly annotated to only 7276 articles. Using propagation to broader descriptors, *Eukaryote* is found to be related to 7.700.235 articles (~87% of all publication in the KG), which seems more realistic according to the expected representation of articles related to Human and model organisms in PubMed. While the co-occurrence increases by 82.293 articles, the association is then not supported by the test (*p* − *value* = 1). The MeSH descriptors annotating the articles bearing the co-occurrences (Supplementary Figure S1) consist of mostly unrelated sets of terms, from *Plant Roots* to *Time Factors*, which also have low importance scores indicating that they are not specifically associated with this relation, but widely distributed in the literature.

#### 3.1.2 New associations inferred from semantic relatedness

An example of an association revealed using propagation is *Oxyfedrine*, a vasodilator and beta-agonist agent, and the disease family of *myocardial ischemia*^13^. Originally, no publication directly links Oxyfedrine to myocardial ischemia: it is only linked to specific diseases of this class: *Angina Pectoris, Coronary Disease* or *Myocardial Infarction*. By propagating annotation to broader terms, the union of articles associated to these diseases supports the relation with *myocardial ischemia*, as well as the higher concepts of *Cardiovascular Diseases* and *Heart Diseases* which are also found to be highly associated with Oxyfedrine. By looking at the other annotations of the supporting corpus (Supplementary Figure S2), we see that unlike the association between Water and Eukaryota, it bears a specific set of topics, with high importance scores, related to Heart Diseases (*Blood Pressure, Coronary Circulation*, etc.).

However, propagation through broader descriptors up to the highest hierarchical concepts, have sometimes yielded reasonable associations of average interest. An example is the association between *Levocabastine*, an anti-histaminic agent used in treatment for conjunctivitis allergies, and the *Eukaryota* kingdom. Their relatedness is completely inferred from semantic relationships as there are no publications explicitly annotated with both. The supporting literature is mainly focused on concepts such as *ophthalmic solutions, anti-allergic agents* and the organisms *Human, Rats* and *Mice* (Supplementary Figure S3), which yield the association of Levocabastin with the whole kingdom.

### 3.2 Relevance of extracted associations

In order to evaluate our approach, we used previously published test-cases from the Metab2MeSH article to determine a reference set of diseases related to Cyclic AMP, and compounds related to Phenylketonuria using literature evidence (data were respectively retrieved from the supplementary Figures 2 and 3 of the original Metab2MeSH article[29]).

#### 3.2.1 Diseases related to Cyclic AMP

Most of the 13 disease-related MeSH associated with Cyclic AMP (cAMP) are malignant neoplasms, such as neuroblastoma or glioma. The top 20 of MeSH descriptors reported by FORUM is compared to the reference set (see Supplementary table S1). FORUM retrieves all diseases annotated in the reference set as over-represented in the cAMP corpus. Also, by taking advantage of the true-path rule and the semantic relations between MeSH terms, it enriches results with new associations involving broader descriptors representing the disease families in which the previously identified disorders belong. For instance, *Neoplasms, Neuroepithelial* is a parent in the MeSH Thesaurus of *Glioma* and *Neuroblastoma*, and *Parathyroid Diseases* is the direct parent of *Hypoparathyroidism* and *Hyperparathyroidism*. This higher level of description allows the direct identification of the type of disorders that could be related to a given compound, in addition to reporting the list of specific diseases.

#### 3.2.2 Compounds related to Phenylketonuria

In the second test case, we compare results for compounds associated with Phenylketonurias (Supplementary tables S2 and S3), a group of metabolic diseases induced by a defect in the production of phenylalanine hydroxylase [43]. Some compounds (12/25) associated with this disease in the reference set are not present in our compound list (in red), as there are no or few articles associated with them in our Knowledge Graph. This could have stemmed from compound disambiguation by PubChem links providers, since some of them were rarely occurring D-configuration amino acids, while L-forms are still found. Differences in the approach used to link literature to compounds, as well as changes in the PubChem database since 2012, make comparison with previous methods difficult. For compounds newly associated with phenylketonuria in our list, they are for the most part pterin derivatives. Supplementary tables S4 and S5 describe results obtained using ChEBI and ClassyFire ontologies providing a semantic description of molecules. This representation highlights molecule families associated with Phenylketonuria rather than simple compounds, like the family of *aromatic amino acids, biopterins and derivatives* or *pterins* whose members were also found significantly associated with the disease in the previous analysis.

#### 3.2.3 Deeper association extraction

We used FORUM to extract significant associations between chemical entities and biomedical concepts and then to explicitly connect them in the KG. This new layer of relations enriches the KG, allowing us to explore and study more complex questions.

We illustrate the potential of our query engine coupled with the extended KG using a previously reported association extracted from a complex chain of relationships. Literature-based discovery was first introduced by Swanson et al. [44] and aimed to extract new hypotheses from initially separated pieces of knowledge developed in scientific articles. In his first example, he proposed as a new hypothesis, that fish oil might be interesting in treatment of Raynaud’s syndrome, by connecting knowledge that a fish oil diet is rich in Eicosapentaenoic acid (EPA), which has antiplatelet actions and increases production of PGI3. Secondly, PGI3 is an analog of PGI2, a vasodilatator and antiplatelet agent, used in treatment of Raynaud’s syndrome. All building blocks for reconstructing Swanson’s reasoning can be found in our KG, using the created *skos:related* relations between entities, from the subgraph shown on Figure 3. The relation between therapeutic MeSH descriptors and the MeSH descriptor representing Raynaud’s syndrome have been extracted using the same approach as described in methods, similarly to [13]. The SPARQL requests that can be used to retrieve these connections are provided in Supplementary documents.

**Figure 3:**
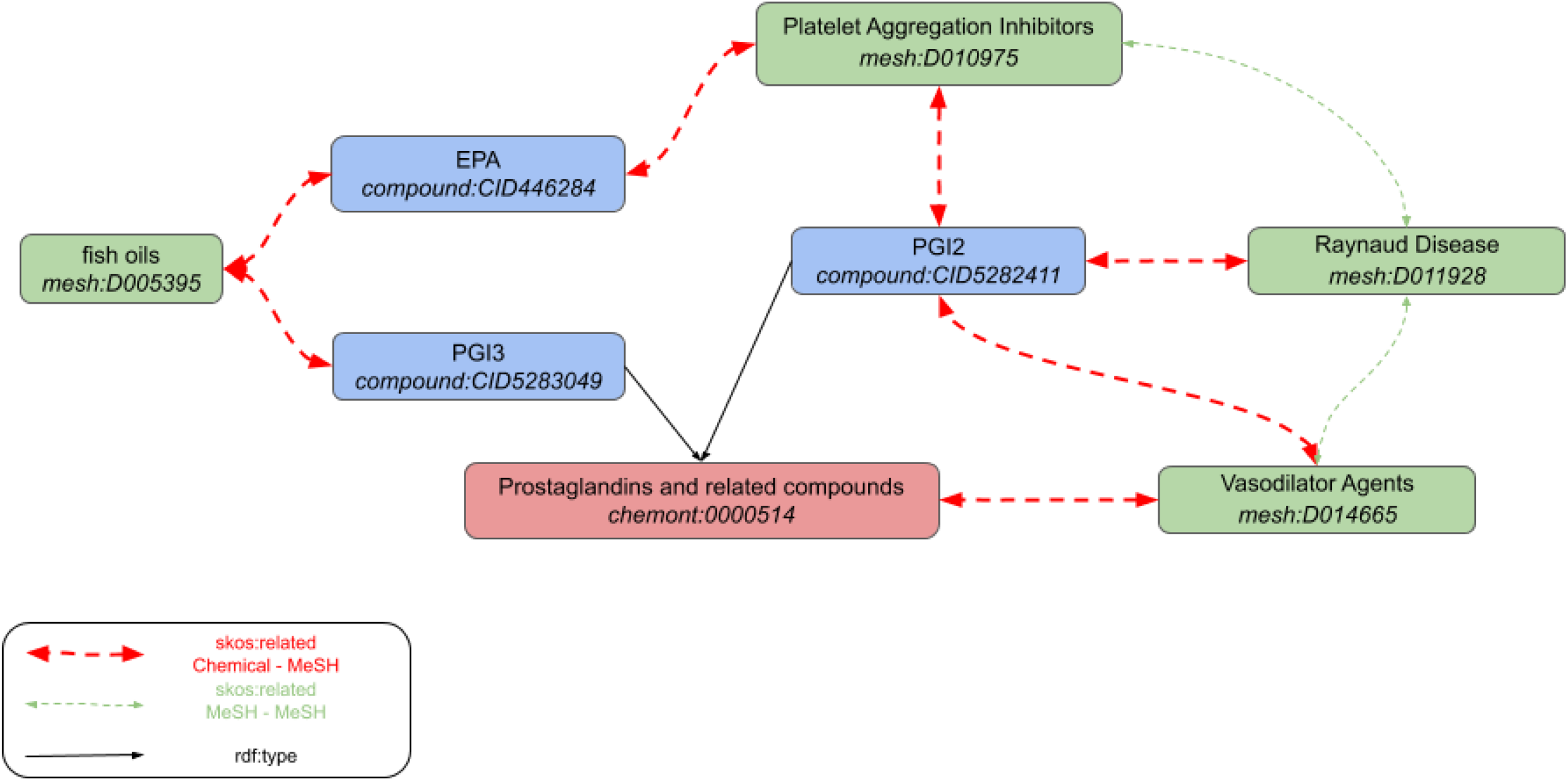
Retrieving the Swanson’s hypotheses fragments in FORUM KG. All skos:related associations have been built from the processing of underlying chains of links found in the federation of linked datasets. There are also *skos:related* links directly linking the chemical class of *Prostaglandins and related compounds* to *fish oils, Platelet Aggregation Inhibitors, Vasodilator Agents* and *Raynaud Disease* in the KG, but those are not shown for clarity reasons.

## 4 Discussion

Federating knowledge is an essential process to succeed in inferring novel hypotheses from metabolomics. For this purpose, we gathered and connected complementary resources in a Knowledge Graph to provide a semantic path between chemical and biomedical concepts through the scientific literature. By performing an enrichment analysis on their related corpora, we estimated the effect-size and relevance of more than 120 million of assertions and selected 8.7 million of them to be instantiated, using a threshold on the resulting q-value at 1*e* − 6. According to this threshold, we statistically expect less than 9 false created relations in the KG, providing a robust resource to explore connections between chemical entities and biomedical concepts.

We have shown that FORUM results are consistent with reported test cases and enriched with new relevant associations, derived from initial outcomes. Metab2MeSH matched chemical compound names to MeSH concepts to determine links to articles, which are, however, a subset of all links that can be extracted using Elink, also taking advantage of external providers and publishers. Moreover, the semantic description of data, combined with the use of the *true-path* rule, allow propagating annotations through the hierarchical relations defined between entities. This enables the computation of associations with several degrees of abstraction from both chemical and biomedical perspectives, allowing to consider disease or chemical families instead of simple compounds or diseases, which can be useful to offer a broader vision on results and guide interpretation. Thus, we have been able to relate such specific disease families like *Neoplasms, Neuroepithelia* and *Parathyroid diseases* to the cyclic AMP and some chemical classes like *pterins* and *aromatic amino acids* to phenylketonuria [43]. Here, we mainly focused our examples on disease cases, but the other MeSH trees are also meaningful and present the similar behavior regarding the impact of the *true-path* rule.

The approach used in FORUM allows us to retrieve twice as many associations than without using the MeSH structure. As MeSH descriptors are organised in trees, the corpora extension is null for leaves and becomes stronger as we get closer to the roots, which therefore gather all articles referring to the main topics. Newly extracted relations are thus associated with MeSH descriptors describing increasingly broader concepts, with larger corpora, enabling the creation of new hypotheses. For example, we were able to significantly relate the Oxyfedrine to the *Myocardial Infarction* disease family. Also, knowing that associated descriptors highlight a drug related context (vasodilator agent and Adrenergic beta agonist), this may suggest hypotheses for the use of Oxyfedrine for other diseases, alternative forms of myocardial ischemia. Since some precise concepts may not always be extracted from relations, abstraction to chemical classes or biomedical broader concepts can still help to find relevant information at a higher level of abstraction. In the same way, we also know that most chemical compounds are not well described in the scientific literature, because for instance, they can’t be correctly identified with current experimental methods. However, for these compounds, concepts that have been linked to the closest chemical class, using articles discussing other members of that class, can provide relevant prior knowledge. The Chemont structural ontology is particularly suited to this purpose as the classification of any molecule can be determined using structural information such as SMILES or InchiKey identifiers. Compounds potentially related to understudied phenotypes could also be estimated by looking at a more broader disease family. For broader descriptors affected by the propagation, it can also help to discard irrelevant relations. Indeed, the co-occurrence between some chemicals and MeSH descriptors may originally appear significant, but mostly because the related MeSH corpus can be underestimated without considering the literature related to sub-concepts (e.g. *Eukaryota*). While the extracted relations can thus be used to identify diseases related to one particular compound for instance, these also represent knowledge fragments that can be assembled to highlight new hypotheses. Indeed, like Swanson’s deduction extracted from our KG, by following paths between relations to infer new connections between concepts and/or chemicals, FORUM could also support knowledge discovery in metabolomics.

### 4.1 Limits

Since FORUM knowledge extraction is based on literature mining, it obviously strongly depends on the quality of the considered corpora. While it can be expected that products of scientific misconduct and malpractice are marginal in the PubMed corpus, it can have a substantial impact on association based on small corpora. For example, one particular harmful practice consisting of multiplying articles from single experiments, coined “salami-slicing” [45, 46, 47], may pass peer-review and evade retractation, while still inducing bias in the contingency table and thus potentially yield spurious associations. Beyond the articles’ content, the metadata quality, namely MeSH indexing and PubChem annotation [18], have also a critical impact on the results. To support the identification of associations that could be yielded by such practices, we propose the *fragility index*, which estimates the minimum number of spurious articles that, if removed, would trigger the exclusion of the association in the KG.

The use of a threshold on the q-value is a convenient and widely used statistical approach to select relevant results from hypothesis testing, but this can also lead to misinterpretation, and those should be treated with care [48]. While statistically relevant associations could be used to derive new hypotheses, on the other hand, the absence of associations shouldn’t be used to this end (Open-world assumption). Indeed, the KG being incomplete by nature and by design, as path-search is at the core of its exploitation, irrelevant shortcuts should be avoided as much as possible. We thus focused on conservative methods to retain ‘strongest’ associations. Other values helping to gauge the effect size and the volatility are provided in complement to q-value.

By increasing the number of associations, including somewhat redundant information with close parent concepts, we also tend to favor information overload. Indeed, we aim at gathering a large collection of strong associations for computational analysis, consequently, the output can result in a large collection of machine-readable associations that can be difficult to navigate manually. Some associations shared by closely related MeSH descriptors, may increase the number of triplets to read without adding much information. Also, associations with overly broad concepts can be highly statistically significant but may not be useful or relevant depending on the context. However, some methods [49] exist to filter such information and have been already applied in similar fields (eg. Gene Ontology), which is a perspective of improvement for FORUM.

The relations drawn between compounds and medical concepts are solely based on non-independence expectation, meaning that a scientific article mentioning the former are more likely to also mention the latter. This doesn’t guarantee any direct relationship, and if one exists, its nature is unknown. Thus, without further inquisition, one should not assume that an association between a compound and a disease imply, for example, its use as a treatment of said disease. Beyond therapeutic uses, the associations can encompass, among others: implication in physiopathology, use as a biomarker for diagnosis, a reagent in related analytical tools, a metabolic product of drug side-effects, and so on. Some text-mining methods could be applied to abstracts of articles supporting significant associations to unveil the nature of the relationship, pushing further the analysis of FORUM results. We have shown that even simple MeSH frequency analysis of association supporting corpus can also provide some insights (See also Imidocarb example in Supplementary Figure S4.).

We have also identified some subtleties related to the propagation process, as associations inferred to broader concepts are not necessarily relevant for all sub-classes. In the example of Levocabastine and Eukaryota, we see that the supporting corpus is mainly related to ophthalmic solutions in human use, but end up relating the compound to a class of organism where presence of conjunctiva is an exception. These types of relations can nonetheless generate new hypotheses but it requires further analyses, and generally associations with broader concepts should be interpreted as “related to SOME x” and not “related to ALL x”. Further developments need to be performed to correctly identify generalization opportunities from inferred relations to broader concepts or chemical classes. Finally, associations with all MeSH categories are included for the sake of exhaustivity and their relevance left at the judgement of the user. We nonetheless believe that the *organisms* category should be interpreted with care, as many large Phylums display associations driven by very few overrepresented model organisms. For example, 81% of articles related to Eukaryota are derived from Human, Rats or Mice studies.

### 4.2 Conclusion

The Knowledge Graph of FORUM provides a resource to connect chemical entities to biomedical concepts through millions of scientific articles, fully accessible and searchable, offering a useful tool to support the interpretation of results in metabolomics and yield new hypotheses. Built with an in-depth annotation of the literature, and collected using approaches from data mining to manual curation, relations are extracted from large corpora analysis, and are therefore less prone to errors than text-mining approaches which require a second examination, as a price for higher descriptiveness. Extracted relations also create higher-level relations whose connections can be explored and easily expanded, through the semantic framework, by linking to larger knowledge collections such as Wikipathway [50] to infer new hypotheses. Indeed, the more data are accessible and linked between various resources, the more we open the path to infer new connections and reveal hidden relations. Moreover, while provided associations only involve one chemical entity regarding a precise biomedical concept, FORUM can also be used to give insights on custom questions. Indeed, when the issue can be formulated as an enrichment problem using a reference set of articles that can be semantically described in the KG, the SPARQL endpoint can be used to extract required data by combining several MeSH descriptors or chemical classes in a request for instance. Interfacing SPARQL requests using some dedicated frameworks such as AskOmics [51] could also improve accessibility and manipulation of the KG.

By gathering knowledge from various resources and inferred assertions inside a FAIR infrastructure, organized around a linked data model, we think that FORUM can be an interesting tool, both to support the interpretation of metabolomics data in several contexts, and to serve as a sand-box to give clues on various questions. As the availability of linked data in life science continues to grow, we hope that FORUM could also be considered as a proof of concept demonstrating the efficiency of this type of approach in information extraction.

## Acknowledgments

The authors express their gratitude to the PubChem, NCBI Eutils and MetaNetX support teams for helping us in the exploitation of their datasets. We thank the Askelys team who provided insight and expertise on the semantic web technology, as well as our colleagues from the EMPREINTE CATI group for their technical support. We are also very grateful to Juliette Cooke for proofreading the manuscript. This project has received funding from the INRA SDN and the European Union’s Horizon 2020 research and innovation program under grant agreement GOLIATH No. 825489.

## Footnotes

1 Ontology and Thesaurus are both structured vocabularies, but thesaurus have a lower power of reasoning than an ontology [11]

2 https://www.ncbi.nlm.nih.gov/books/NBK25500/

3 ftp://ftp.ebi.ac.uk/pub/databases/chebi/ontology/chebi.owl

4 http://classyfire.wishartlab.com/downloads

5 https://protege.stanford.edu/

6 ftp://ftp.ncbi.nlm.nih.gov/pubchem/RDF/

7 https://dataguide.nlm.nih.gov/eutilities/utilities.html

8 https://www.research.ibm.com/labs/almaden/index.shtml

9 ftp://ftp.nlm.nih.gov/online/mesh/rdf/

10 DCMI namespace for the Dublin Core metadata element set. https://www.dublincore.org/

11 SKOS Simple Knowledge Organization System https://www.w3.org/TR/skos-reference/

12 https://hub.docker.com/r/tenforce/virtuoso

13 A disorder of cardiac function caused by insufficient blood flow to the muscle tissue of the heart.

