## Supplementary materials for "FORUM: Building a Knowledge Graph from public databases and scientific literature to extract associations between chemicals and diseases"

#### S1 Supplementary tables

| RANK | MESH_LABEL | MESH | PubMed Articles | q-value | odds-ratio | chisq-stat |
|---|---|---|---|---|---|---|
| 1 | Pseudohypoparathyroidism | D011547 | 234 | 1.30e-277 | 48.38 | 7749 |
| 2 | Neuroblastoma | D009447 | 662 | 1.02e-260 | 5.76 | 2463 |
| 3 | Neuroectodermal Tumors, Primitive, Peripheral | D018241 | 666 | 1.50e-260 | 5.72 | 2452 |
| 4 | Neuroectodermal Tumors, Primitive | D018242 | 675 | 1.56e-239 | 5.15 | 2141 |
| 5 | Neoplasms, Neuroepithelial | D018302 | 1082 | 1.17e-184 | 2.86 | 1258 |
| 6 | Neoplasms, Germ Cell and Embryonal | D009373 | 1786 | 8.33e-171 | 2.13 | 1026 |
| 7 | Neuroectodermal Tumors | D017599 | 1637 | 8.07e-163 | 2.17 | 985 |
| 8 | Neoplasms, Nerve Tissue | D009380 | 1662 | 1.43e-159 | 2.13 | 959 |
| 9 | Cystic Fibrosis | D003550 | 365 | 4.90e-98 | 3.95 | 772 |
| 10 | Leydig Cell Tumor | D007984 | 114 | 1.04e-94 | 18.54 | 1628 |
| 11 | Sertoli-Leydig Cell Tumor | D018310 | 117 | 1.51e-86 | 14.60 | 1311 |
| 12 | Parathyroid Diseases | D010279 | 429 | 4.44e-82 | 3.06 | 576 |
| 13 | Cholera | D002771 | 125 | 8,62e-80 | 11.17 | 1050 |
| 14 | Graves Disease | D006111 | 249 | 3,97e-77 | 4.48 | 643 |
| 15 | Calcium Metabolism Disorders | D002128 | 606 | 3.05e-74 | 2.37 | 466 |
| 16 | Sex Cord-Gonadal Stromal Tumors | D018312 | 126 | 1.40e-72 | 9.46 | 875 |
| 17 | Neoplasms, Gonadal Tissue | D018309 | 126 | 2.55e-71 | 9.20 | 848 |
| 18 | Exophthalmos | D005094 | 249 | 5.34e-69 | 4.05 | 548 |
| 19 | Polycystic Kidney Diseases | D007690 | 133 | 3.11e-64 | 7.34 | 680 |
| 20 | Adrenal Gland Neoplasms | D000310 | 293 | 1.18e-63 | 3.34 | 463 |
| 23 | Glioma | D005910 | 552 | 2.16e-59 | 2.23 | 363 |
| 26 | Hypoparathyroidism | D007011 | 126 | 1.11e-52 | 6.12 | 508 |
| 27 | Hyperparathyroidism | D006961 | 307 | 2.29e-51 | 2.80 | 345 |
| 32 | Cell Transformation, Neoplastic | D002471 | 459 | 6.72e-46 | 2.15 | 276 |
| 34 | Hypercalcemia | D006934 | 207 | 5.31e-42 | 3.17 | 297 |
| 38 | Adrenal Cortex Neoplasms | D000306 | 109 | 2.44e-38 | 5.04 | 335 |
| 42 | Osteosarcoma | D012516 | 199 | 8.89e-33 | 2.77 | 217 |

Table S1: Top 20 hits of FORUM search for compound "Cyclic AMP", only considering disease descriptors. Lines in grey represent diseases which are in the reference set but which are ranked below the top 20 in this list. Lines in green represent newly associated MeSH descriptors and those in white are descriptors of the reference set also found in our top 20.

| RANK | Metabolite | CID | PubMed Articles | P-Value adj | FoldChange | chisq-Stat |
|---|---|---|---|---|---|---|
| 1 | L-phenylalanine | 6140 | 2279 | 0 | 222.1 | 499735.3 |
| 2 | endophenyl | 6925665 | 2279 | 0 | 222.1 | 499735.3 |
| 3 | D-phenylalanine | 71567 | 2279 | 0 | 222.1 | 499735.3 |
| | DL-Phenylalanine | 994 | 2279 | 0 | 219.8 | 499735.0 |
| 5 | Sapropterin | 44257 | 278 | 0 | 296.4 | 80864.7 |
| 6 | AC1L9H5R | 444951 | 278 | 0 | 296.4 | 80864.7 |
| 7 | 6,7-Dihydrobiopterin | 133246 | 278 | 0 | 296.2 | 80918.0 |
| 8 | 7-tetrahydrobiopterin | 1125 | 278 | 0 | 295.9 | 80824.7 |
| 9 | 1-(2-amino-4-hydroxy-5,6,7,8-tetrahydropteridin-7-yl) propane-1,2-diol | 169715 | 278 | 0 | 295.9 | 80824.7 |
| 10 | Pterin H B2 | 2380 | 353 | 0 | 206.0 | 71398.0 |
| 11 | biopterin | 444475 | 353 | 0 | 206.0 | 71398.0 |
| 12 | 7,8-Dihydrobiopterin | 252 | 353 | 0 | 200.3 | 69422.2 |
| 13 | L-tyrosine | 6057 | 538 | 0 | 28.0 | 14032.7 |
| 14 | D-Tyrosine | 71098 | 538 | 0 | 28.0 | 14032.7 |
| 15 | DL-Tyrosine | 1153 | 538 | 0 | 28.0 | 14032.7 |
| 16 | 2-amino-3-(4-hydroxyphenyl)propanote | 5460807 | 538 | 0 | 28.0 | 14032.7 |
| 17 | (2S)-2-Azaniumyl-3-(4-hydroxyphenyl)propanoate | 6942100 | 538 | 0 | 28.0 | 14032.7 |
| 18 | (2S)-2-amino-3-(4-hydroxyphenyl)propanoate | 5460822 | 538 | 0 | 28.0 | 14032.7 |
| 19 | (2R)-2-amino-3-(4-hydroxyphenyl)propanoate | 5460814 | 538 | 0 | 28.0 | 14032.7 |
| 20 | LS-188017 | 24848110 | 542 | 0 | 24.3 | 12107.4 |
| 21 | Phenylpyruvic acid | 997 | 39 | 2.07e-85 | 339.4 | 13159.0 |
| 22 | Enol-phenylpyruvate | 641637 | 39 | 2.07e-85 | 339.4 | 13159.0 |
| 23 | 2-hydroxy-3-phenylprop-2-enoic acid | 691 | 39 | 2.07e-85 | 339.4 | 13159.0 |
| 24 | Tryptophan | 1148 | 120 | 3.72e-67 | 8.9 | 841.6 |
| 25 | L-Tryptophan-beta-14C | 148495 | 120 | 3.72e-67 | 8.9 | 841.6 |

Table S2: Reference set of compounds related to phenylketonurias, from the Top 25 hits of Metab2MeSH search. Lines in red represent compounds which are not found using our knowledge base.

| RANK | Metabolite | CID | PubMed Articles | q-value | odds-ratio | chisq-stat |
|---|---|---|---|---|---|---|
| 1 | (2S)-2-Azaniumyl-3-phenylpropanoate | 6925665 | 2744 | 0 | 899.6 | 717859.4 |
| 2 | Phenylalanine | 6140 | 3045 | 0 | 949.4 | 695241.7 |
| 3 | 2-Amino-6-[(1S,2R)-1,2-dihydroxypropyl]-5,6,7,8-tetrahydropteridine-4(1H)-one | 136153088 | 436 | 0 | 567.0 | 178498.9 |
| 4 | Tetrahydrobiopterin | 135402045 (eq. of 1125) | 436 | 0 | 567.0 | 178498.9 |
| 5 | 5,6,7,8-Tetrahydrobiopterin | 135409384 | 436 | 0 | 567.0 | 178498.9 |
| 6 | Sapropterin dihydrochloride | 135409471 | 436 | 0 | 567.0 | 178498.9 |
| 7 | Trihydroxybutyrophenone | 129630809 | 436 | 0 | 567.0 | 178498.9 |
| 8 | Sapropterin | 135398654 (eq. of 44257) | 436 | 0 | 566.6 | 178420.0 |
| 9 | Tetrahydrodictyopterin | 135433600 | 436 | 0 | 566.3 | 178341.3 |
| 10 | 2,4,5-Trihydroxybutyrophenone | 15008 | 436 | 0 | 564.0 | 177791.9 |
| 11 | D-Erythro-Biopterin | 135449517 | 517 | 0 | 390.9 | 150772.5 |
| 12 | Orinapterin | 135738580 | 517 | 0 | 390.9 | 150772.5 |
| 13 | Biopterin | 135403659 (eq. of 444475) | 517 | 0 | 390.9 | 150772.5 |
| 14 | d-Threo biopterin | 135909519 | 517 | 0 | 390.9 | 150772.5 |
| 15 | Pterin H B2 | 135398729 (eq. of 2380) | 521 | 0 | 381.0 | 148652.0 |
| 16 | 4(1H)-Pteridinone,2-amino-7-(1,2-dihydroxypropyl)-5,6,7,8-tetrahydro- | 135616732 (eq. of 169715) | 195 | 0 | 558.4 | 82872.6 |
| 17 | (6S)-2-Amino-6-[(1S,2S)-1,2-dihydroxypropyl]-5,6,7,8-tetrahydro-3H-pteridin-4-one | 136003108 | 184 | 0 | 361.1 | 54034.9 |
| 18 | L-Tyrosinate(1-) | 5460822 | 597 | 0 | 35.6 | 16907.2 |
| 19 | (2S)-2-Azaniumyl-3-(4-hydroxyphenyl)propanoate | 6942100 | 597 | 0 | 35.6 | 16907.2 |
| 20 | L-Tyrosine | 6057 | 654 | 0 | 16.2 | 7797.9 |
| 21 | Phenylpyruvic acid | 997 | 62 | 1.33e-128 | 350.5 | 18030.0 |
| 22 | Sodium phenylpyruvate monohydrate | 23666336 | 57 | 2.26e-127 | 519.6 | 23312.9 |
| 23 | Sodium phenylpyruvate | 23667645 | 57 | 2.26e-127 | 519.6 | 23312.9 |
| 24 | 2-Oxo-3-phenylpropanoate | 4592697 | 57 | 2.26e-127 | 519.6 | 23312.9 |
| 25 | L-Tryptophan | 6305 | 141 | 2.87e-72 | 7.9 | 807.6 |
| 30 | DL-Tryptophan | 1148 | 124 | 4.23e-70 | 9.0 | 846.8 |
| 58 | Dihydrobiopterin | 135398687 (eq. of 252) | 11 | 4,09e-20 | 184.8 | 1684,4 |

Table S3: Top 25 hits of FORUM search for disease Phenylketonurias (MeSH D010661). Lines in grey represent compounds which are in the reference set but which are ranked lower in this list. Lines in green represent newly associated compounds and those in white are compounds of the reference set also found in our list. Some entries in the Metab2MeSH results correspond to old PubChem identifiers, so we looked for equivalences in current PubChem entries. We annotated them with *eq. of*.

| RANK | CHEBI LABEL | CHEBI ID | PubMed Articles | q.value | odds-ratio | chisq-stat |
|---|---|---|---|---|---|---|
| 1 | phenylalanine | CHEBI:28044 | 3045 | 0 | 996.4 | 638213.3 |
| 2 | 5,6,7,8-tetrahydrobiopterin | CHEBI:15372 | 436 | 0 | 518.7 | 163713.7 |
| 3 | tetrahydropterin | CHEBI:30436 | 440 | 0 | 485.6 | 156910.2 |
| 4 | biopterin | CHEBI:15373 | 521 | 0 | 350.7 | 136557.8 |
| 5 | biopterins | CHEBI:22881 | 524 | 0 | 342.6 | 134150.8 |
| 6 | erythrose 4 - phosphate / phospho-enolpyruvate family amino acid | CHEBI:73690 | 3210 | 0 | 204.5 | 125691.2 |
| 7 | aromatic amino acid | CHEBI:33856 | 3289 | 0 | 154.1 | 87093.1 |
| 8 | amino acid zwitterion | CHEBI:35238 | 2930 | 0 | 110.0 | 83289.9 |
| 9 | proteinogenic amino acid | CHEBI:83813 | 3336 | 0 | 61.7 | 32802.9 |
| 10 | L-alpha-amino acid | CHEBI:15705 | 3383 | 0 | 56.0 | 27991.1 |
| 11 | alpha-amino acid | CHEBI:33704 | 3404 | 0 | 51.7 | 25055.2 |
| 12 | tyrosinate(1-) | CHEBI:32784 | 597 | 0 | 32.9 | 15455.9 |
| 13 | tyrosine | CHEBI:18186 | 654 | 0 | 15.0 | 7078.0 |
| 14 | L-alpha-amino acid anion | CHEBI:59814 | 629 | 0 | 14.5 | 6616.0 |
| 15 | alpha-amino-acid anion | CHEBI:33558 | 629 | 0 | 14.5 | 6582.9 |
| 16 | amino-acid anion | CHEBI:37022 | 629 | 0 | 14.5 | 6581.2 |
| 17 | pterins | CHEBI:26375 | 563 | 0 | 15.2 | 6344.7 |
| 18 | pteridines | CHEBI:26373 | 563 | 0 | 14.9 | 6194.7 |
| 19 | diol | CHEBI:23824 | 441 | 7.03e-312 | 13.5 | 4518.9 |
| 20 | polar amino acid | CHEBI:26167 | 894 | 5.02e-249 | 4.4 | 1846.7 |

Table S4: Top 20 hits of FORUM search for disease Phenylketonurias (MeSH D010661) using propagation through the ChEBI ontology.

| RANK | CHEMONT LABEL | CHEMONT ID | PubMed Articles | q.value | odds-ratio | Chisq_stat |
|---|---|---|---|---|---|---|
| 1 | Biopterins and derivatives | C0001651 | 553 | 0 | 250.3 | 107357.0 |
| 2 | Pterins and derivatives | C0000110 | 593 | 0 | 16.6 | 7375.3 |
| 3 | Tyrosine and derivatives | C0004319 | 718 | 0 | 14.1 | 7177.0 |
| 4 | Pteridines and derivatives | C0000109 | 600 | 0 | 14.5 | 6394.8 |
| 5 | Indolyl carboxylic acids and derivatives | C0001290 | 207 | 1.47e-110 | 8.4 | 1268.4 |
| 6 | Phenylpyruvic acid derivatives | C0001276 | 62 | 2.08e-110 | 169.8 | 9321.2 |
| 7 | Serotonins | C0001637 | 158 | 1.64e-36 | 3.5 | 265.5 |
| 8 | Histidine and derivatives | C0004311 | 103 | 8.07e-36 | 4.9 | 309.7 |
| 9 | Tryptamines and derivatives | C0000183 | 159 | 2.06e-34 | 3.3 | 244.4 |
| 10 | Indole-3-acetic acid derivatives | C0001252 | 45 | 5.64e-23 | 7.6 | 246.1 |
| 11 | Leucine and derivatives | C0004329 | 71 | 1.42e-17 | 3.6 | 127.4 |
| 12 | 2(hydroxyphenyl)acetic acids | C0004644 | 12 | 1.34e-14 | 41.1 | 420.8 |
| 13 | Phenylacetic acids | C0000418 | 12 | 1.60e-14 | 40.5 | 414.0 |
| 14 | Phenylpropanoic acids | C0002551 | 45 | 2.77e-14 | 4.4 | 112.6 |
| 15 | Methionine and derivatives | C0004143 | 59 | 3.21e-14 | 3.5 | 100.6 |
| 16 | Pyridoxines | C0001948 | 28 | 7.95e-14 | 7.1 | 138.7 |
| 17 | Isoleucine and derivatives | C0004330 | 26 | 2.62e-11 | 6.0 | 102.2 |
| 18 | Catecholamines and derivatives | C0000182 | 99 | 4.0e-08 | 1.9 | 42.6 |
| 19 | D-alpha-amino acids | C0004145 | 33 | 2.55e-06 | 2.9 | 37.7 |
| 20 | L-cysteine-S-conjugates | C0004555 | 19 | 2.97e-06 | 4.3 | 44.6 |

Table S5: Top 20 hits of FORUM search for disease Phenylketonurias (MeSH D010661) using propagation through the ClassyFire (ChemOnt) ontology. The last two lines are not significant at the considered threshold.

### S2 Supplementary Figures

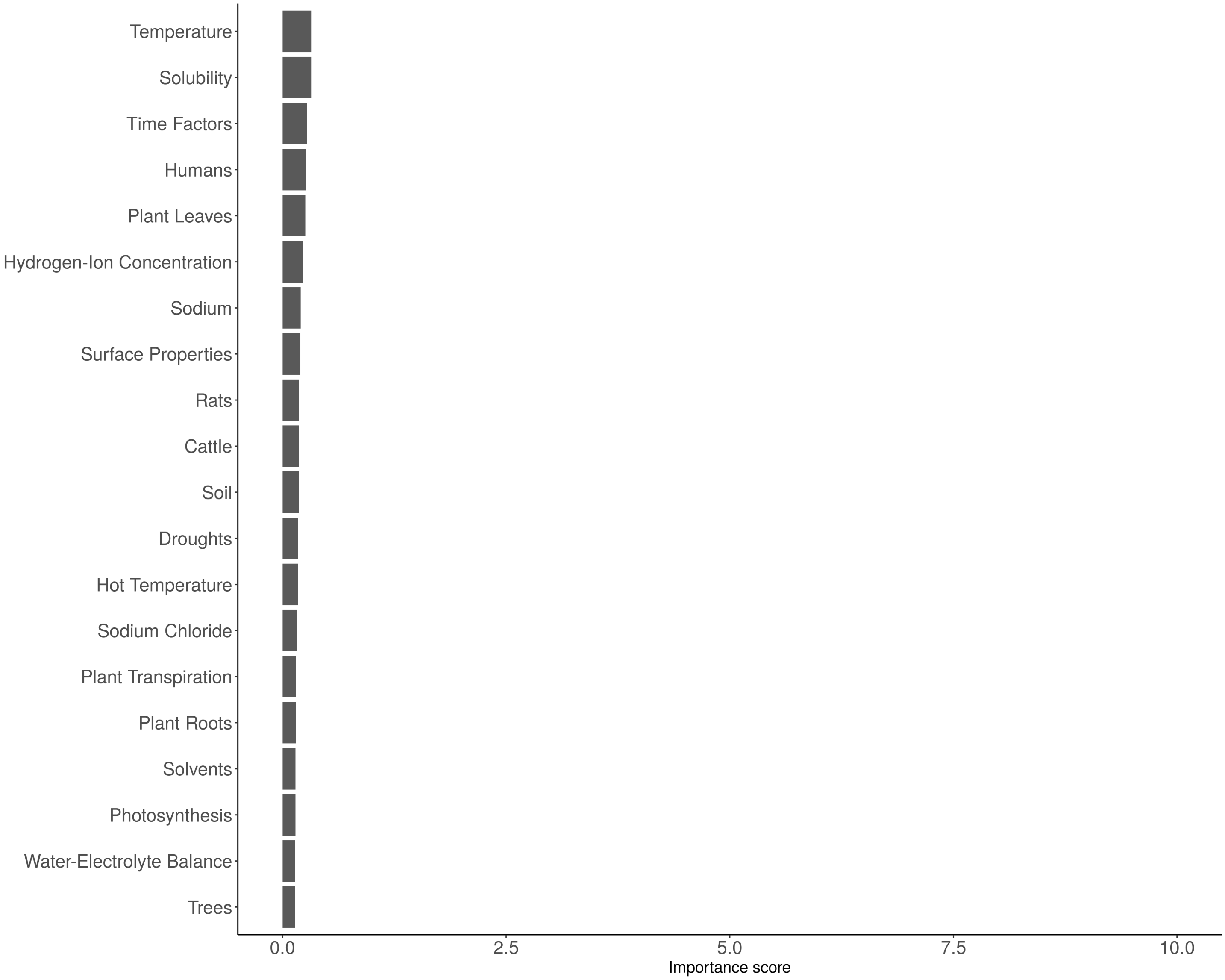


Figure S1: Top 20 most important descriptors describing the relation between Water and Eukaryote. The MeSH descriptor associated with Water (*mesh:D014867*) was removed from the selection as it directly represents the studied compound.

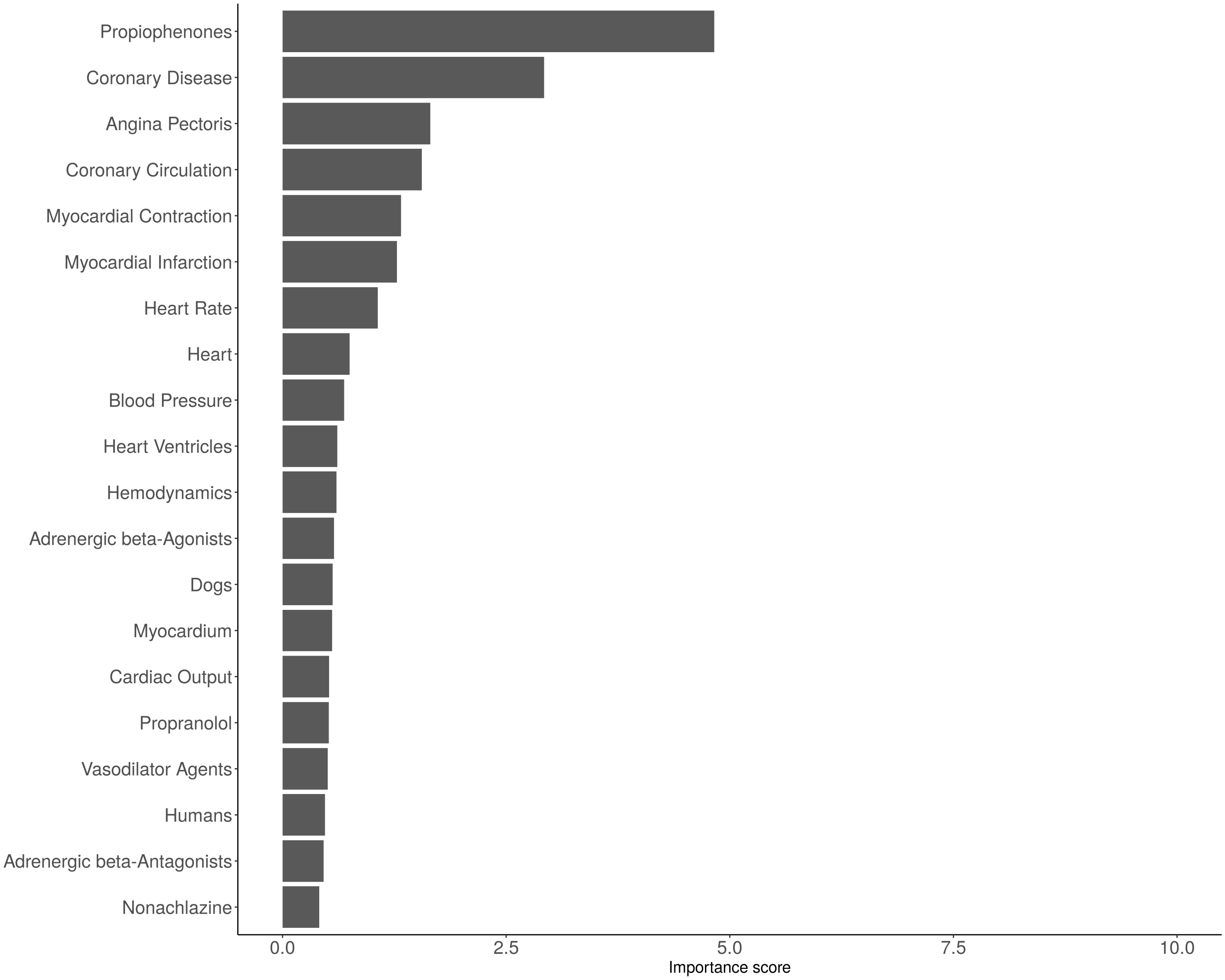


Figure S2: Top 20 most important descriptors describing the relation between Oxyfedrine and Myocardial Ischemia. The MeSH descriptor associated with Oxyfedrine (*mesh:D010099*) was removed from the selection as it directly represents the studied compound.

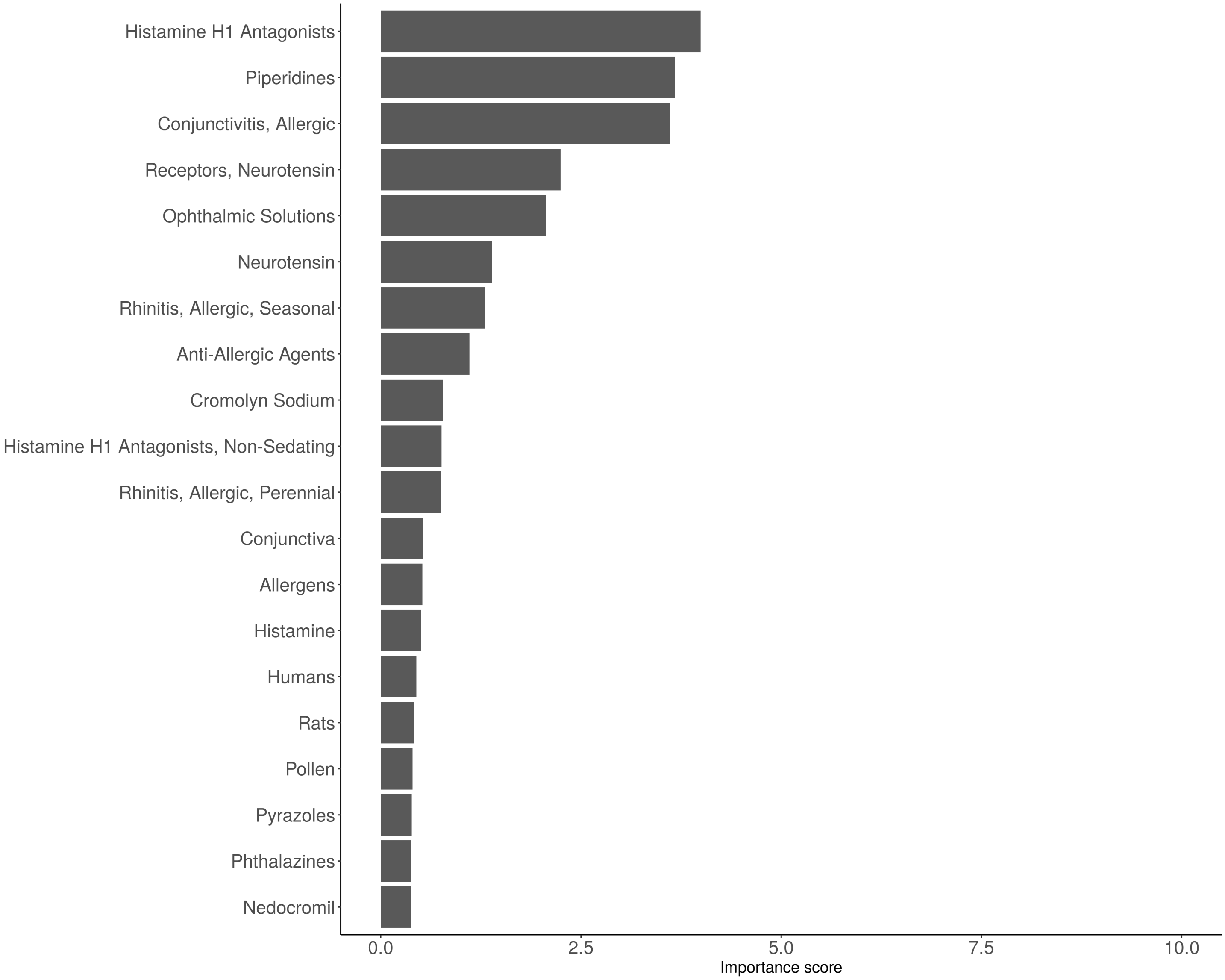


Figure S3: Top 20 most important descriptors describing the relation between Levocabastine and Eukaryote.

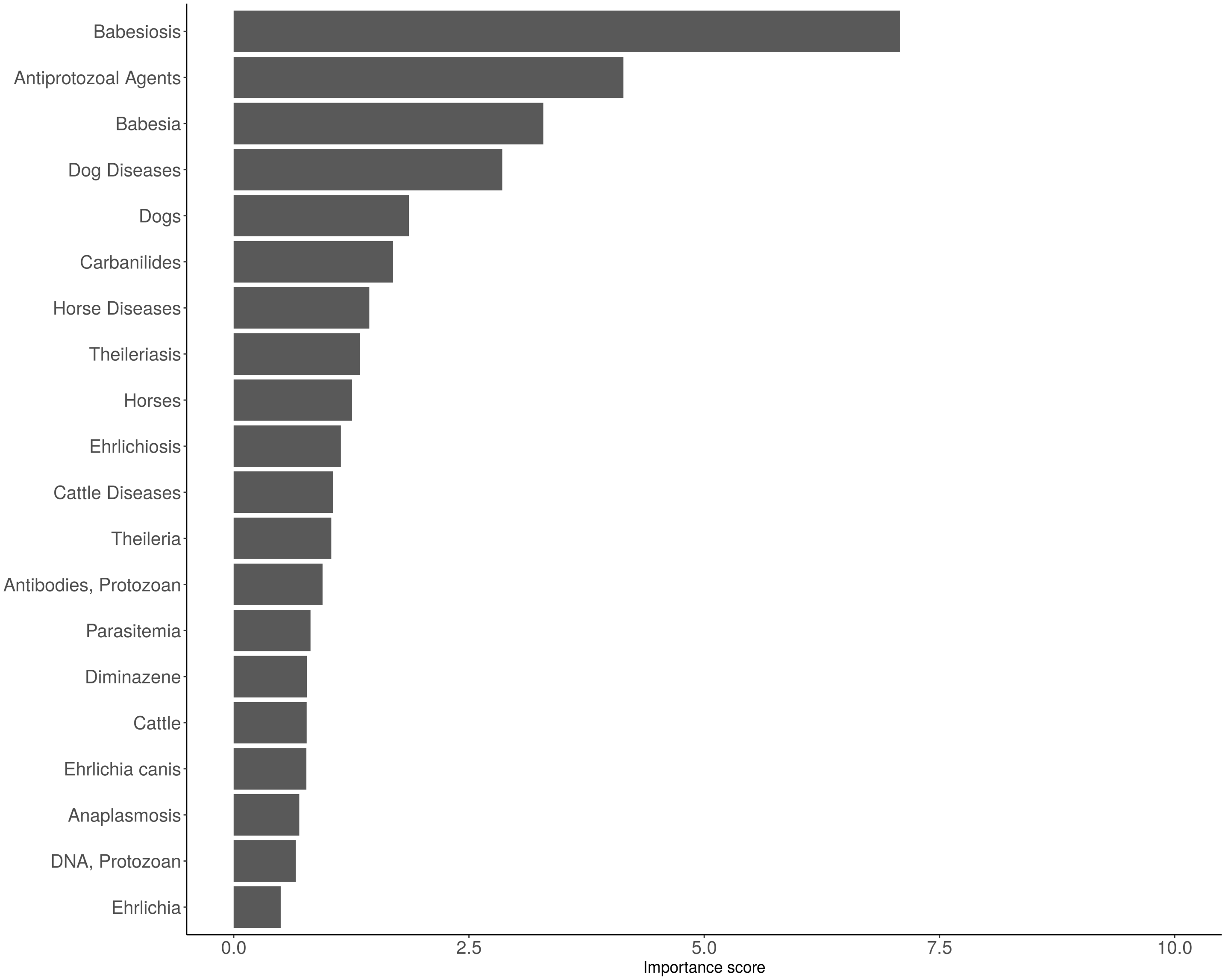


Figure S4: Top 20 most important descriptors describing the relation between Imidocarb and Tick diseases. The MeSH descriptor associated with Imidocarb (*mesh:D007095*) was removed from the selection as it directly represents the studied compound.

### S3 Supplementary material

#### S3.1 Supplementary Results: Impact of the semantic level on association extraction

##### S3.1.1 True-path rule impact on MeSH corpora size

The true-path rule modifies corpora sizes and thus co-occurrence counts used for statistical testing of independence. The impact of the true-path rule depends on the vocabulary structure, as a "flat" ontology would bring few changes. It also depends on the indexation practice, since overuse of the broadest descriptor would also lead to small changes. In order to explore the overall impact of the semantic level integration to our knowledge network, we checked how it influenced the corpus of MeSH descriptors and their extracted relations with PubChem compounds.

We first looked at the delta in corpus size associated with each MeSH descriptor when propagating, or not, the annotations through the MeSH Thesaurus, according to the *true-path* rule (Figure S5). We chose to organise MeSH in categories by number of childs, to observe the relation between the tree location and the potential benefit of the use of semantic relations. MeSH descriptors with no childs ([0-1[) can be considered as leaves in the MeSH Tree, and the use of semantic relations has no impact on their corpus size, as there are no more specific terms in the Thesaurus from which to propagate the annotations. Benefits in terms of the number of added publications increase with the hierarchical position of MeSH descriptors in the tree.

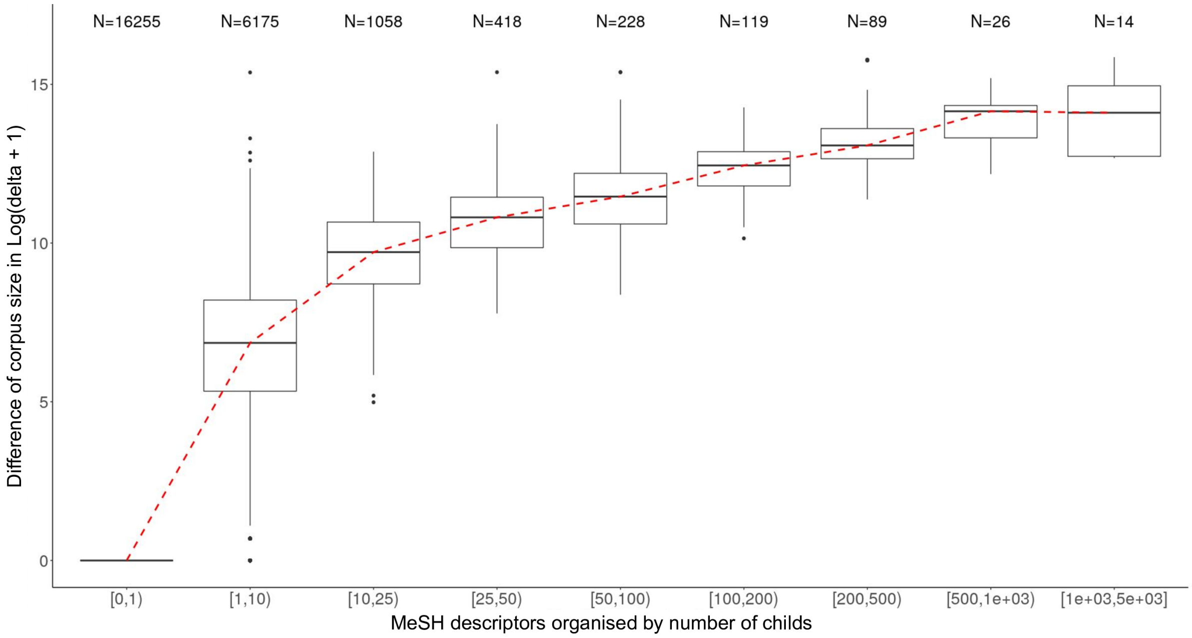


Figure S5: Comparison of the difference in corpus size associated with 24.382 MeSH descriptors with and without propagation through the MeSH Thesaurus. 117 MeSH originally have an empty corpus without propagation. MeSH are organised by number of children according to the MeSH tree, from leaves with no children [0-1[, like Gardner Syndrome (*mesh:D005736*), to root descriptors like Neoplasm (*mesh:D009369*). For each category, the number of MeSH descriptors that belong to it is indicated at the top of the boxplot. The red dashed line indicates the mean.

##### S3.1.2 True-path rule impact on association broadness

The MeSH corpus size distribution was compared between associations gained, discarded and kept (Figure S7). Corpus sizes were determined using the number of articles annotated to each MeSH descriptor after propagation according to the *true-path rule*. The *conserved* set is composed of 22.051 distinct MeSH descriptors ($\sim$ 97% of all studied MeSH with significant associations) and the *added* and *lost* sets are respectively composed of 7.622 and 7.769 MeSH descriptors. The distributions of MeSH corpus size are very similar for these last two as they share a lot of descriptors in common. However, these descriptors are not specific to added or lost associations as they are also involved in conserved relations (Cf. Figure S6). Only 11 MeSH concepts aren't associated with any compounds after applying the true-path rule, such as: *Civilisation (D002962)* or *Archives (D001109)*. Also, 625 were added without having any associated compounds initially, like *Toxic Actions* (D004786). Finally, 59 MeSH are found both in the newly added associations and the lost ones, without being found in the conserved set, meaning that their associated compounds have completely changed. The case *Adnexal Diseases (D000291)* is an example of such changes. This term was originally rarely used for annotation (172 articles), resulting in few significant associations with compounds (only 2). However, its corpus has been heavily impacted by the true-path rule propagation (42.051 new articles added), since its narrower child concepts were much more used. This has diluted previous relations and revealed new ones from the new corpus.

Furthermore, there is a gap between the distribution of MeSH involved in conserved associations and those involved in added or lost associations (Cf. Figure S7). It seems that by using propagation, affected associations implies a subset of broader MeSH descriptors whose corpus are on average larger than those linked to conserved associations.

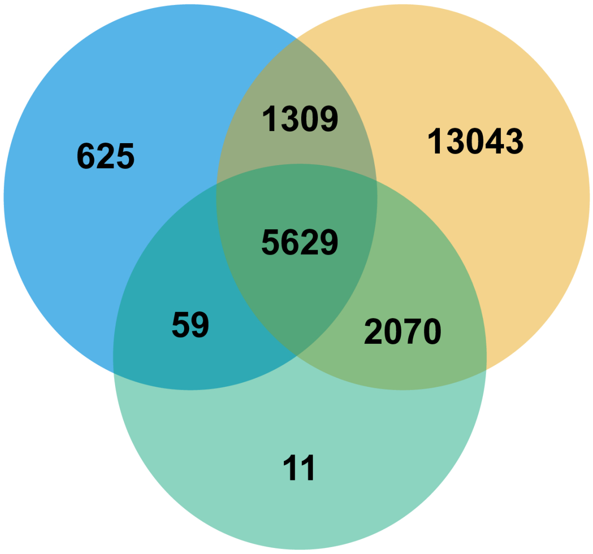


Figure S6: A Venn diagram of MeSH sets associated with conserved (orange), lost(green) or added (blue) associations. Among the 24.382 studied MeSH descriptors, 22.746 are involved in significant associations with PubChem compounds.

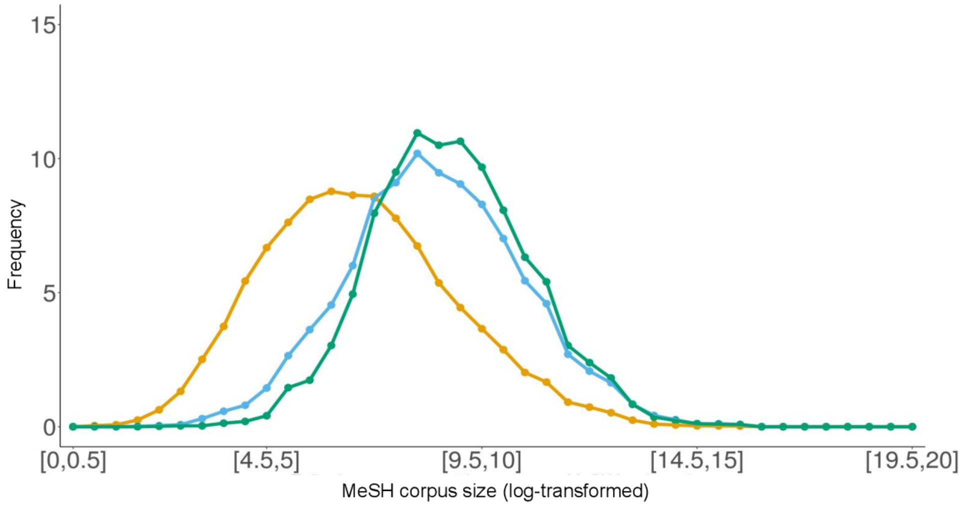


Figure S7: Corpus size distribution (Log-transformed) of MeSH sets associated with each category of the V enn diagram. The Corpus size of MeSH descriptors was determined with literature propagation through the MeSH Thesaurus.

##### S3.1.3 True-path rule impact on MeSH representation in associations

The impact of applying ontology based reasoning on extracted associations has been characterized considering MeSH tree positions. We first checked the distribution of significant associations among the MeSH trees by considering whether they are new, conserved or lost, Figure S8. For all MeSH categories, selected associations are in similar proportion compared to the whole set observed on the venn diagram ($\sim$ 50% of added, $\sim$ 47% conserved and $\sim$ 3% lost), except for the organisms category for which 63% of associations are new. See details on Supplementary table S6. Similar distributions are also found for subclasses of the Diseases MeSH tree (Figure S8). Differences in the absolute number of associations in each MeSH tree are directly linked to their global rate of annotation in the KG. As all our studied articles are retrieved from PubChem entry mentions, they might be more likely to be related with chemical compounds or activities, thus the *Chemicals & Drugs* tree is the most represented in our results ($\sim$ 52%). MeSH descriptors from *Infections* and *Neoplasm* sub-trees are likewise among the most annotated disease categories in PubMed.

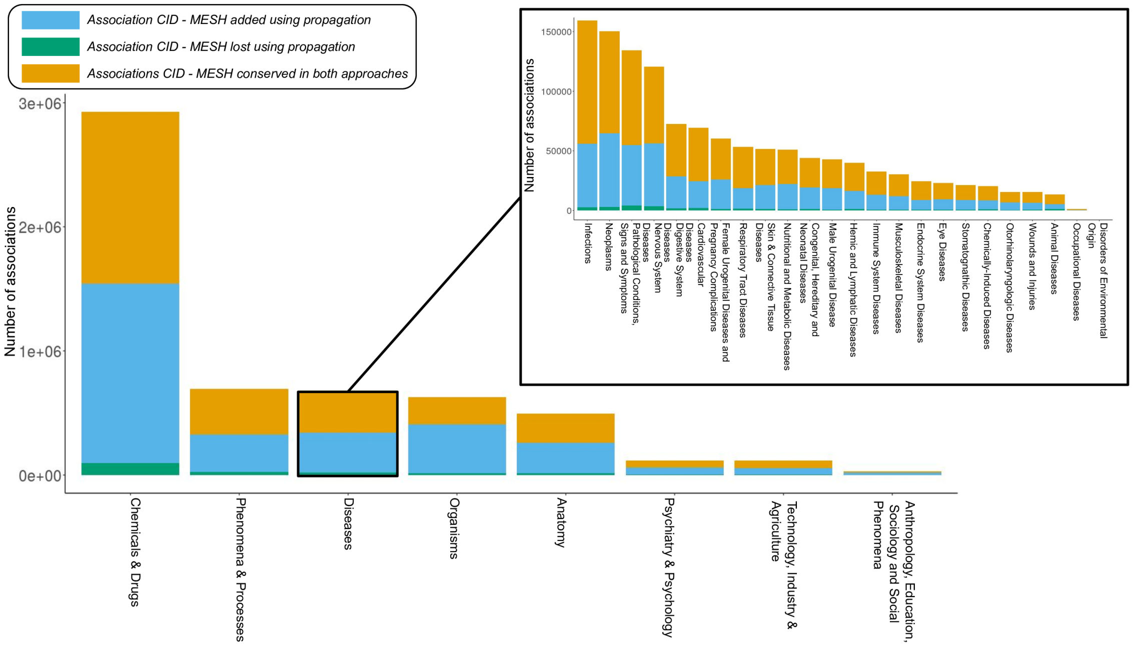


Figure S8: Stacked barplot describing the number of associations CID - MESH involved in each MeSH tree and organised by categories of the venn diagram (Cf. Figure 2). The distribution of associations in the Disease tree is also described in detail.

| Tree | N, Added | N, Lost | N, Conserved | % Added | % Lost | % Conserved |
|---|---|---|---|---|---|---|
| Chemicals and Drugs | 1446010 | 100705 | 1381315 | 49.38 | 3.44 | 47.17 |
| Phenomena and Processes | 301684 | 23912 | 366490 | 43.59 | 3.46 | 52.95 |
| Diseases | 320130 | 20475 | 337174 | 47.23 | 3.02 | 49.75 |
| Organisms | 393382 | 12547 | 220060 | 62.84 | 2.00 | 35.15 |
| Anatomy | 243582 | 14078 | 238380 | 49.11 | 2.84 | 48.01 |
| Psychiatry and Psychology | 53540 | 2886 | 59716 | 46.1 | 2.48 | 51.41 |
| Technology, Industry, and Agriculture | 58032 | 4143 | 54035 | 49.93 | 3.57 | 46.50 |
| Anthropology, Education, Sociology, and Social Phenomena | 16790 | 962 | 13663 | 53.45 | 3.06 | 43.50 |
| **Global** | **2772905** | **173704** | **2581608** | **50.16** | **3.14** | **46.70** |

Table S6: Number of added, lost and conserved associations with related percentages by MeSH trees. **Warning:** As one MeSH descriptor can be located in several different trees, counts for the Venn diagram do not have to be interpreted as the total.

#### S3.2 Supplementary Methods

##### S3.2.1 Fragility index

The *fragility index* aims at determining the number $n$ of articles that, if removed from the corpus, would return an unsignificant *p-value*. To determine $n$, we estimate several scenarios in which a growing number of articles supporting the relation between for instance, a compound (or a chemical class) $A$ and a MeSH descriptor $B$ would be removed. Nonetheless, not all scenarios can be tested for computational reasons and we need to establish bounds in which to determine scenarios. We used the Jeffrey interval (at 95%) to determine a confidence interval around the proportion of articles discussing a MeSH descriptor $B$ among those discussing the compound $A$, $\frac{N_{ij}}{N_{i.}}$ with $N_{ij}$ the number of articles discussing the MeSH $j$ and the compound $i$ (the co-occurrences) and $N_{i.}$ the total number of articles discussing the compound $i$. We compute this proportion regarding the compound corpus size, because most of them are unfortunately not well described in the literature, contrary to studied MeSH descriptors. Also, the Jeffrey interval is particularly suited to determine confidence intervals on a small sample size [42].

Using the lowest bound $p_{min}$ of the interval, we estimate the co-occurrence by rounding $N_{min}$, corresponding to the scenario that led to such minimal proportion: $N_{min} \approx p_{min}N$.

We first test using this lowest scenario: if the test is still significant, the association is declared robust and no other tests are computed, else, we compute all possible scenarios from $N_{min}$ to the observed co-occurrence, to determine the first scenario that fails. Using this failing scenario, we determine $n$, the number of articles that if removed from the corpus, makes the *p-value* over the significance threshold ($1e-6$). The *q-value* being always higher or equal than the *p-value*, this also informs us about the *q-value* that could be obtained from this scenario. See examples in Figures S9 and S10.

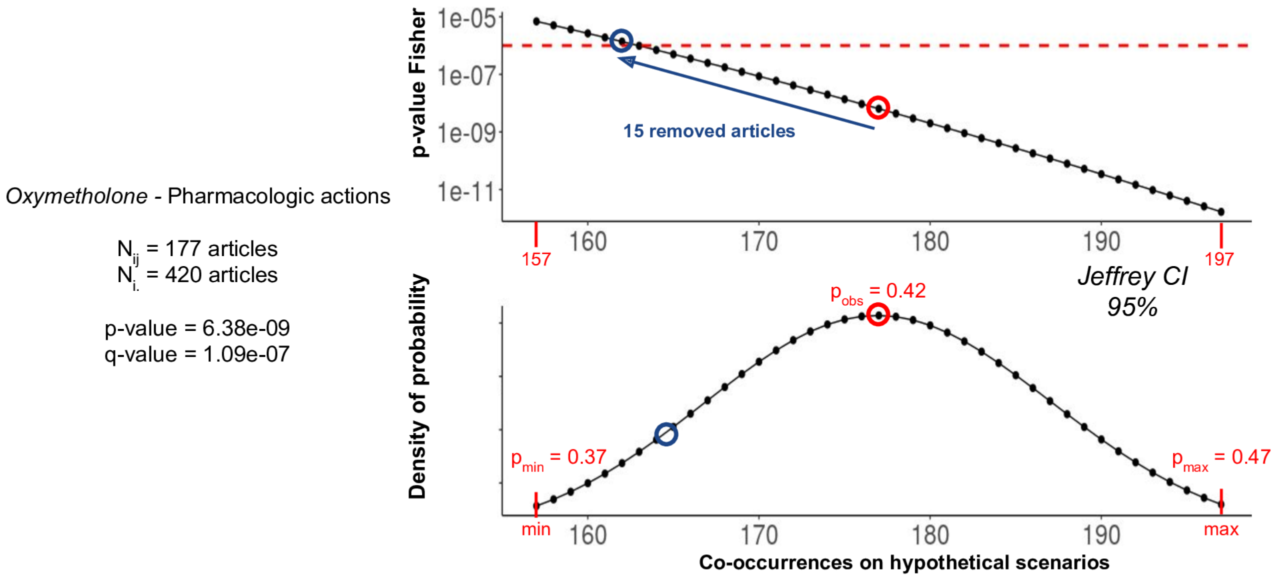


Figure S9: Example Oxymethone - Pharmacologic actions of the fragility index procedure

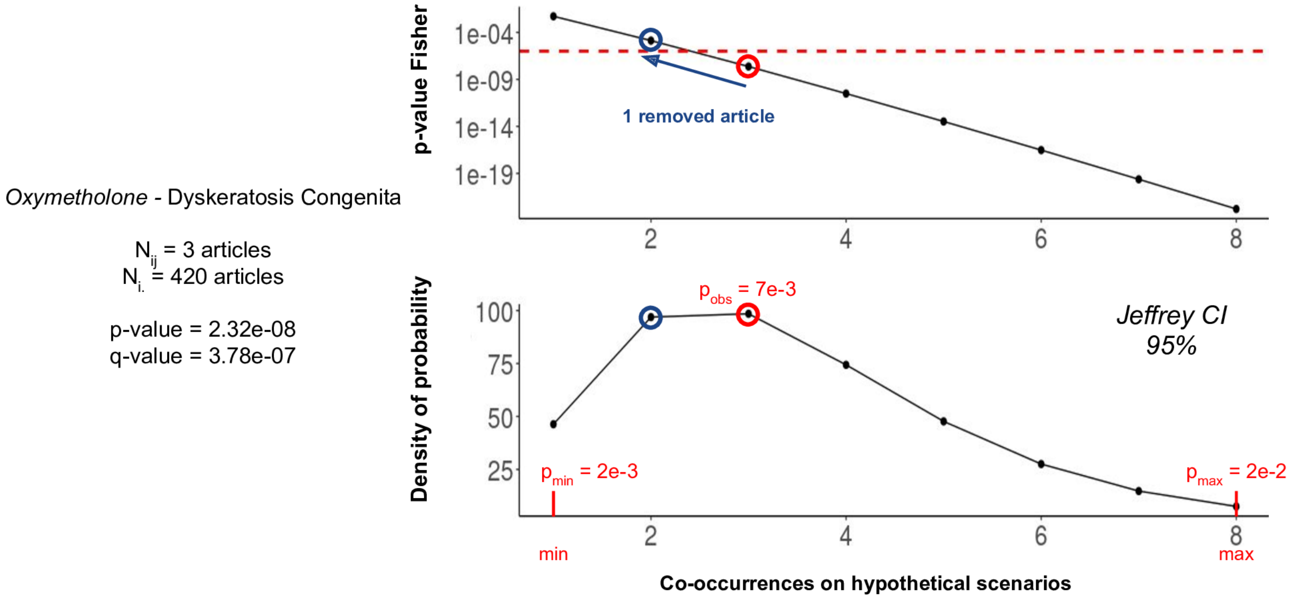


Figure S10: Example Oxymethone - Dyskeratosis Congenita of the fragility index procedure

The association between *Oxymethone* (an androgen and anabolic steroid) and *Pharmacologic actions* is supported by 177 articles, among 420 related to Oxymethone, corresponding to a proportion of $\approx 0.42$. *Pharmacologic actions* is a broad descriptor with about 2.5 million articles. Using the lower bound of the Jeffrey CI on this proportion, we estimated the co-occurrence at the lowest scenario: 157 articles. Then, we computed each independence test between the lowest (157 articles) and the observed scenario (177 articles). From the obtained *p-values*, it appears that 15 articles should be removed from the association corpus to obtain a *p-value* higher than the considered threshold (and therefore also the q-value), and thus could change the decision on the relation relevance. The interpretation of this result according to the corpus size of the compound, remains to the user, but in this case, the limit of 15 articles seems to be sufficiently high to be confident about this relation.

In a second example, we studied the relation between *Oxymethone* and *Dyskeratosis Congenita* (a X-linked recessive syndrome). This relation is only supported by 3 articles, but is nonetheless significant, notably because *Dyskeratosis Congenita* is rarely discussed in the KG (111 articles). Following the same procedure, we found that just by removing one article, this relation could no longer be significant. As there are only three supporting articles, it could be important to check their relevance in order to validate this relation.

The *Fragility index* is complementary to the *q-value* by giving a more formal interpretation of the strength of a relation, and, as shown on previous examples, it can also highlight the impact of the corpus size. Indeed, due to the discrete nature of the data, when studying rare MeSH descriptors or compounds, therefore having small corpora, small variations in the co-occurence can have a significant impact on statistical tests performed.

##### S3.2.2 Ranking MeSH descriptors supporting a relation using a TF-IDF score

To estimate the importance of each MeSH descriptor supporting a relation, we propose to compute a score, analog to the TF-IDF, using the frequency of the MeSH descriptor in the corpus of articles supporting the relation, compared to its frequency in the whole KG.

So, to estimate the importance of a MeSH descriptor $k$ annotated in publications supporting the relation between a compound $i$ and a MeSH descriptor $j$:

$$Score = \frac{N_{i,j}^{k}}{N_{i,j}} \times log(\frac{N_{.k}}{N_{..}})$$

- $N_{i,j}$ the number of articles supporting the co-occurrence between the compound $i$ and the MeSH descriptor $j$.
- $N_{i,j}^{k}$ the number of articles discussing the MeSH descriptor $k$ among those supporting the relation between $i$ and $j$.
- $N_{.k}$ The total number of articles discussing $k$ in the KG.
- $N_{..}$ The total number of articles in the KG.

#### S3.3 Supplementary data: Swanson's Hypothesis - SPARQL requests

Here, we propose two SPARQL requests that can be used to retrieve links involved in Swanson's reasoning.

```
DEFINE input:inference "schema-inference-rules"
PREFIX rdf: <http://www.w3.org/1999/02/22-rdf-syntax-ns#>
PREFIX rdfs: <http://www.w3.org/2000/01/rdf-schema#>
PREFIX xsd: <http://www.w3.org/2001/XMLSchema#>
PREFIX owl: <http://www.w3.org/2002/07/owl#>
PREFIX meshv: <http://id.nlm.nih.gov/mesh/vocab#>
PREFIX mesh: <http://id.nlm.nih.gov/mesh/>
PREFIX voc: <http://myorg.com/voc/doc#>
PREFIX cito: <http://purl.org/spar/cito/>
PREFIX fabio:    <http://purl.org/spar/fabio/>
PREFIX owl: <http://www.w3.org/2002/07/owl#>
PREFIX void: <http://rdfs.org/ns/void#>
PREFIX cid:   <http://rdf.ncbi.nlm.nih.gov/pubchem/compound/>
PREFIX sio: <http://semanticscience.org/resource/>
PREFIX obo: <http://purl.obolibrary.org/obo/>
PREFIX skos: <http://www.w3.org/2004/02/skos/core#>
```

```
prefix dcterms: <http://purl.org/dc/terms/>
PREFIX chemont: <http://purl.obolibrary.org/obo/CHEMONTID_>

select distinct (strafter(STR(?other_compound),"http://rdf.ncbi.nlm.nih.gov/pubchem/compound/CID")
as ?CID)

from <https://forum.semantic-metabolomics.org/ClassyFire/direct-parent/2020>
from <https://forum.semantic-metabolomics.org/EnrichmentAnalysis/CID_MESH/2020>
from <https://forum.semantic-metabolomics.org/ChemOnt/2016-08-27>
from <https://forum.semantic-metabolomics.org/MeSHRDF/2020-12-07>
where
{
    {
        select ?other_compound
        where
        {
            mesh:D011928 skos:related ?RD_c .
            ?RD_c skos:related ?mesh_TU .
            ?mesh_TU (meshv:treeNumber|meshv:treeNumber/meshv:parentTreeNumber+) ?tn .
            ?mesh_selected meshv:treeNumber ?tn .
            VALUES ?mesh_selected { mesh:D002317 mesh:D006401 }
            ?other_compound skos:related ?mesh_TU .
            ?other_compound skos:related mesh:D005395 .
            ?other_compound a chemont:0003909 .
            }
        }
    FILTER NOT EXISTS {?other_compound skos:related mesh:D011928}
}
```

In this requests, we extracted:

- Compounds related to the Raynaud's Disease (mesh:D011928)
- MeSH descriptors significantly related to these compounds, by selecting those that are related to therapeutic actions: Cardiovascular Agents or Hematologic Agents.
- We then retrieve other compounds related to these MeSH descriptors
- We also specify that these new compounds must :
  - be related to fish oils (mesh:D005395)
  - belong to the chemical class of Fatty Acyls (chemont:0003909).
  - not be already related to Raynaud's Disease in our KG.

Therefore, these Fatty Acyls compounds associated with fish oils are related to the same therapeutic effects (Cardiovascular Agents or Hematologic Agents) as some compounds used as treatment for Raynaud's Disease. Here we retrieve for instance Eicosapentaenoic acid (EPA), which indeed have the same therapeutic effect (Platelet Aggregation Inhibitors) as PGE1 or PGI2 which are used to treat Raynaud's disease.

```
DEFINE input:inference "schema-inference-rules"
PREFIX rdf: <http://www.w3.org/1999/02/22-rdf-syntax-ns#>
PREFIX rdfs: <http://www.w3.org/2000/01/rdf-schema#>
PREFIX xsd: <http://www.w3.org/2001/XMLSchema#>
PREFIX owl: <http://www.w3.org/2002/07/owl#>
PREFIX meshv: <http://id.nlm.nih.gov/mesh/vocab#>
PREFIX mesh: <http://id.nlm.nih.gov/mesh/>
PREFIX voc: <http://myorg.com/voc/doc#>
PREFIX cito: <http://purl.org/spar/cito/>
PREFIX fabio: <http://purl.org/spar/fabio/>
```

```
PREFIX owl: <http://www.w3.org/2002/07/owl#>
PREFIX void: <http://rdfs.org/ns/void#>
PREFIX cid: <http://rdf.ncbi.nlm.nih.gov/pubchem/compound/>
PREFIX sio: <http://semanticscience.org/resource/>
PREFIX obo: <http://purl.obolibrary.org/obo/>
PREFIX skos: <http://www.w3.org/2004/02/skos/core#>
prefix dcterms: <http://purl.org/dc/terms/>
PREFIX chemont: <http://purl.obolibrary.org/obo/CHEMONTID_>

select distinct (strafter(STR(?other_compound),"http://rdf.ncbi.nlm.nih.gov/pubchem/compound/CID")
as ?CID)
from <https://forum.semantic-metabolomics.org/EnrichmentAnalysis/CID_MESH/2020>
from <https://forum.semantic-metabolomics.org/MeSHRDF/2020-12-07>
where
{
        {
                select distinct ?other_compound
                from <https://forum.semantic-metabolomics.org/EnrichmentAnalysis/CID_MESH/2020>
                from <https://forum.semantic-metabolomics.org/ClassyFire/direct-parent/2020>
                where
                {
                        {
                                select ?RD_class
                                from <https://forum.semantic-metabolomics.org/EnrichmentAnalysis/
                                CHEMONT_MESH/2020>
                                from <https://forum.semantic-metabolomics.org/MeSHRDF/2020-12-07>
                                where
                                {
                                        mesh:D011928 skos:related ?RD_class .
                                        ?RD_class skos:related ?mesh_TU .
                                        ?mesh_TU (meshv:treeNumber|meshv:treeNumber/
                                        meshv:parentTreeNumber+) ?tn .
                                        ?mesh_selected meshv:treeNumber ?tn .
                                        VALUES ?mesh_selected { mesh:D002317 mesh:D006401 }
                                }
                        }
                ?other_compound a ?RD_class .
                ?other_compound skos:related mesh:D005395 .
                FILTER NOT EXISTS {?other_compound skos:related mesh:D011928}
                }
        }
        FILTER NOT EXISTS {
                ?other_compound skos:related ?mesh_TU .
                ?mesh_selected meshv:treeNumber ?tn .
                VALUES ?mesh_selected { mesh:D002317 mesh:D006401 }
                ?mesh_TU (meshv:treeNumber|meshv:treeNumber/meshv:parentTreeNumber+) ?tn
        }

}
```

In this request, we extracted:

- Chemical classes related to Raynaud's Disease which are also related to therapeutic actions: Cardiovascular Agents or Hematologic Agents
- We then selected other members of these chemical classes that are related to fish oils but not to Raynaud's Disease.

- We also specify these compounds must **not** be associated with therapeutic effects (Cardiovascular Agents or Hematologic Agents) in our KG.

These are therefore compounds associated with fish oils that belong to the same chemical class as Raynaud's disease-related compounds with therapeutic effects (cardiovascular agents or haematological agents), but for which no such effects are reported. This allows us to retrieve for instance PGI3, the analog of PGI2, which is used as treatment of the Raynaud's diseases for its properties of vasodilator and antiplatelet agents.
